# Limb connective tissue is organized in a continuum of promiscuous fibroblast identities during development

**DOI:** 10.1101/2024.03.18.585505

**Authors:** Estelle Hirsinger, Cédrine Blavet, Marie-Ange Bonnin, Léa Bellenger, Tarek Gharsalli, Delphine Duprez

## Abstract

Tight developmental coordination of muscle attachments with skeletal muscle is fundamental for building functional limbs and allowing locomotion. Muscle attachments include connective tissue fibroblasts of tendon and muscle connective tissue. Although muscle attachments play critical roles in development, our understanding of connective tissue fibroblast developmental programs lags behind that of other components of the musculoskeletal system, mainly because fibroblasts are highly heterogeneous and poorly characterized. Combining single-cell RNA-sequencing-based strategies including trajectory inference and *in situ* hybridization analyses, we address the diversity of connective tissue fibroblasts and their developmental trajectories during chicken limb fetal development. We show that fibroblasts switch from a program providing positional information to a program of lineage diversification at the onset of fetal period. Muscle connective tissue and tendon contain several fibroblast populations that emerge asynchronously. Once the final muscle pattern is set, *in silico* fibroblast populations that are close in transcriptional identity are found in neighbouring locations in limbs, prefiguring the adult fibroblast layers associated to muscle. We propose that the limb connective tissue is organised in a continuum of promiscuous fibroblast identities, allowing for the robust and efficient connection of the muscular system to bone and skin.

**Highlights:** Dissection of connective tissue heterogeneity using single-cell RNA-sequencing and *in situ* RNA staining

*In silico* fibroblast populations map to distinct fibroblast layers in limbs

Molecular signatures of fetal fibroblast populations at the origin of limb adult connective tissue

Epimysium and endomysium fibroblasts are transcriptionally distinct at fetal stage

Limb fibroblasts display promiscuous identities reflecting their interconnected 3D-network

Fibroblasts switch from providing positional information to differentiation during development

## Introduction

Although connective tissue (CT) plays critical roles in limb development, our understanding of CT fibroblast differentiation lags behind that of other components of the musculoskeletal system mainly because CT fibroblast populations are highly heterogeneous and poorly characterized.

In Vertebrate limbs, CT corresponds to an interconnected three-dimensional network of fibroblasts and extra-cellular matrix (ECM) that support the function of the musculo-skeletal system. Muscle attachments include muscle CT (MCT) and tendon (Helmbacher and Stricker, 2020; Sefton and Kardon, 2019; Nassari et al., 2017). The whole musculoskeletal system is attached to the skin via the hypodermis that is not considered to be part of the dermis *per se* (Lynch and Watt et al., 2018). MCT includes the successive fibroblast layers, epimysium, perimysium and endomysium that surround individual muscles, muscle fiber bundles and muscle fibers, respectively (Purslow et al., 2020). Tendon is described as a continuum of MCT linking muscle to bone (Purslow et al., 2020). Tendon is surrounded by a thin fibroblast layer, the epitenon, then by a loose fibroblast sheath, the peritenon. Both are morphologically distinct from the tendon proper. Paratenon includes peritenon and epitenon layers in adult tissues (Benjamin et al., 2008; Nourissat et al., 2015). MCT and tendon fibroblasts differ by their matrix composition and spatial organization of collagen fibers. Tendons contain collagen fibers displaying a regular organization parallel to the tendon axis, while MCT display collagen fibers with less specific organisation (reviewed in Nassari et al., 2017). The different matrix compositions of each limb CT prefigure different functions and mechanical properties. In contrast to the muscle lineage, there is no identified master gene driving the differentiation programs towards dermis, MCT and tendon, but recognized markers have been identified for each CT type, providing us with robust molecular tools to follow each limb CT type. The bHLH transcription factor TWIST2 (DERMO-1) is expressed in developing dermis (Li et al., 1995) and has been shown to be sufficient to launch the developmental program leading to skin appendage formation in chicken embryos (Hornik et al., 2005). The bHLH transcription factor Scleraxis (SCX) is a recognized marker of developing tendons, and *Scx* mutant mice show tendon defects (Murchinson et al., 2007). The Zinc finger transcription factor OSR1 (Odd-Skipped related-1) identifies a population of developmental fibro-adipogenic progenitors (FAPs) regulating myogenesis (Vallecillo-Garcia et al., 2017) and controlling the pro-regenerative response of FAPs during muscle regeneration (Kotsaris et al., 2023, Stumm et al., 2018).

Limb CT fibroblasts originate from the lateral plate mesoderm, while myogenic cells are mainly derived from somitic mesoderm (Chevallier et al., 1977, Christ et al., 1977). CT is recognized as an important source of extrinsic cues that regulate skeletal muscle differentiation, growth and patterning (reviewed in Duprez 2002; Kardon, 2011; Nassari et al., 2017; Sefton and Kardon, 2019; Helmbacher and Stricker 2020). *Osr1* loss-of*-*function leads to reduced myogenic cell proliferation and survival resulting in limb muscle patterning defects (Vallecillo García et al., 2017). The ablation of tendon cells leads to muscle patterning defect in chicken limbs (Kardon, 1998) and genetic ablation of *Scx*+ tendon cells to alteration of muscle shapes and attachment sites in mouse limbs (Ono et al., 2023). Moreover, *Osr1* gain- and loss-of-function experiments in chicken mesenchymal limb cells showed that OSR1 favors the MCT differentiation program (regulating the expression of *COL3A1* and *COL6A1* genes) at the expense of the tendon differentiation program (Stricker et al., 2012). In addition to OSR1, other transcription factors produced by CT fibroblasts have been shown to regulate limb muscle patterning during development, such as Tcf4 (now Tcfl2), belonging to the TCF/LEF family (Kardon et al., 2003), the T-box genes Tbx3 and Tbx5 (Hasson et al., 2010, Colasanto et al., 2016), and the homeobox Hoxa11 (Swinehart et al., 2013). However, Tcf4 and Hoxa11 are also expressed in muscle cells (Murphy et al., 2011, Asfour et al., 2023) or in muscle lineage (Flynn et al., 2023), where they could directly act on muscle patterning. It has been recently shown that a subpopulation of CT fibroblasts integrates muscle fibres at the muscle/tendon interface during development and postnatal stages (Esteves de Lima et al., 2021, Yaseen et al., 2021, Yan et al., 2022, Flynn et al., 2023). This unexpected fibroblastic origin of myonuclei at muscle tips close to tendon provides us with a cellular mechanism mediating fibroblast-driven muscle patterning.

Transcriptomic strategies at the level of the cell/nucleus have been used to highlight fibroblast heterogeneity of muscle attachments, providing us with a plethora of CT fibroblast markers in adult tissues. Adult MCT fibroblasts, known as FAPs, are recognized to be highly heterogeneous. Since the first cartography at the cell level of muscle resident cells (Gordani et al., 2019), numerous FAP populations have been identified but with different markers depending on the studies, making it difficult to draw a global picture of the identity and functions of FAP populations (reviewed in Guliani et al., 2022, Fukada and Uezemi, 2023). However, one common feature across these transcriptomic studies is the identification of FAP populations based on low and high levels of molecular markers, such as SCA1 and PDGFRalpha (Guliani et al., 2021, 2022). Although the hierarchical fibroblast layers surrounding muscle are well identified, the FAP populations described *in silico* were not assigned to any of these layers *in situ*.

Similarly to MCT fibroblasts, two to ten tendon fibroblast populations have been identified in adult tendons using transcriptomic analysis of mouse, human or rat tendons (De Micheli et al., 2020, Kendal et al., 2020, Fu et al., 2023, Steffen et al., 2023), including tendon fibroblasts with progenitor properties (Yin et al., 2016, Harvey et al., 2019). No universal molecular signature for these *in silico* fibroblast populations emerged from these studies. However, cells of paratenon and tendon proper show distinct molecular signatures and cellular properties (Dyment et al., 2013; Mienaltowski et al., 2014, 2019; Pechanec and Mienaltowski, 2023).

The studies above addressed CT biology in the adult. To date, there is no data concerning fibroblast populations of muscle attachments during development. In this study, we identified CT fibroblast populations, their molecular signatures and developmental trajectories during chicken limb development and growth using single-cell RNA-sequencing and inferred trajectory analyses. With extensive in situ hybridization analysis, we localized these *in silico* fibroblast populations in fetal limbs and found that they prefigure the mature layers of CT fibroblasts of the musculo-skeletal system.

## RESULTS

### The Dermis, Tendon and MCT branches emerge successively

To exhaustively address fibroblast diversity, we performed single-cell RNA-sequencing (scRNAseq) analysis on whole chicken forelimbs at successive developmental stages, ranging from E4, the progenitor stage, to E10, the final fibro-muscular pattern. We took advantage of the previously-published E4, E6 and E10 datasets (Esteves de Lima et al., 2021) and generated the E7 and E9 datasets following the same protocol. Clustering analysis on E4, E6, E7, E9 and E10 datasets show that CT fibroblasts, identified with known CT markers (*TWIST2*, *OSR1* and *SCX*), segregate from muscle, vessels, blood cells, ectoderm and neural crest cells at all stages (Figure S1A,B for E7 and E9, see Esteves de Lima et al., 2021 for E4, E6 and E10). To identify fibroblast populations within the limb CT, we bio-informatically extracted CT clusters from the whole limb datasets at each developmental stage and performed unsupervised sub-clustering of these CT datasets (Figure S1C). Plotting the distribution of the cell cycle phases across each CT dataset showed that the proportion of proliferating cells (G2M+S phases) drops from 61% at E4 to 42% at E6 and then stabilizes around 35% up to E10 (Figure S1D,E). The five CT datasets were then combined to generate an integrated 5CT dataset containing 24570 cells (Figures 1A, S1F).

**Figure 1.**
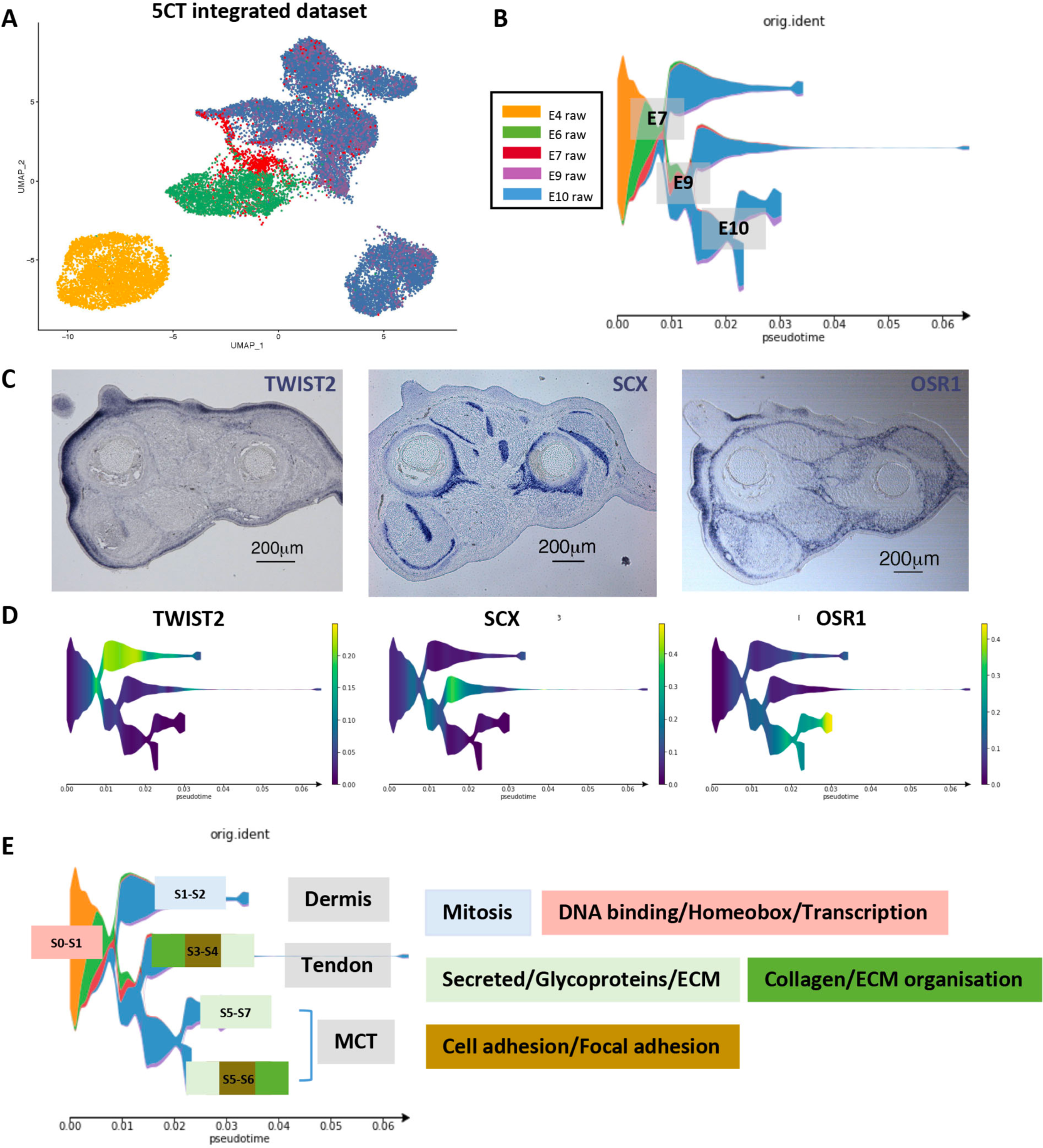
The Dermis, Tendon and MCT branches emerge successively (**A**) UMAP plot of the integrated 5CT dataset (24 570 cells in total), showing the distribution of the CT datasets of origin. (**B**) Dendrogram of the STREAM-derived inferred trajectory for the 5CT dataset, showing the distribution of fibroblasts according to their CT datasets of origin. **(C)** Colorimetric *in situ* hybridization to adjacent and transverse limb sections of E10 chicken embryos with TWIST2, OSR1 and SCX probes (blue staining). **(D)** Dendrogram plotting gene expression levels for *TWIST2*, *OSR1* and *SCX* on the inferred trajectory of the 5CT dataset. **(E)** Dendrogram as in (B) showing the association of color-coded DAVID categories with the origin (S0-S1), Dermis (S1-S2), Tendon (S3-S4) and MCT (S5-S7 and S5- S6) branches. DAVID category-associated GO terms were grouped as follows: DNA binding, Homeobox, Transcription regulation in pink; Secreted, Glycoproteins and Extra Cellular Matrix in light green; Collagens, Extra Cellular Matrix organization, Fibrillar collagen in green; Cell adhesion, Focal adhesion, Proteoglycan in brown. For the S3-S4 and S5-S6 branches, the top 3 associated DAVID categories are ordered by decreasing enrichment scores with the highest score to the left. The other branches are associated with a single DAVID category.

In order to visualize the emergence of major CT types during development, trajectory inference using STREAM (Chen et al., 2019) was performed on the integrated 5CT dataset (Figure 1A,B). Three major branches arise successively (Figure 1B). To identify each branch, we plotted the expression of markers for the main CT types, *i.e* TWIST2 for dermis (Li et al., 1995), OSR1 for MCT (Vallecillo-García et al, 2017) and SCX for tendon (Murchinson et al., 2007) (Figure 1C,D). Dermis was the first CT type to emerge at E7, followed by Tendon and MCT around E8/E9. MCT subsequently split into two branches at E10 (Figure 1B-D). We conclude that the Dermal, Tendon and MCT types emerge successively during limb development.

### CT fibroblasts switch from a positional information to a lineage diversification program at the onset of the fetal period

To identify the main biological activities associated with each branch, we performed a functional enrichment analysis on genes identified by STREAM (Table S1) as having branch-specific expression using DAVID (Sherman et al., 2022, Huang et al. 2009). Categories associated with GO terms such as “DNA binding“, “Homeobox” and “Transcription regulation” were over-represented in the S0-S1 origin branch. Except for the Dermis branch (S1-S2) that was associated with mitosis-related categories, the Tendon (S3-S4) and MCT (S5-S7, S5-S6) branches were associated with ECM-related categories (Figure 1E). The Tendon branch and to a lesser extent the S5-S6 MCT branch exhibited more mature categories of ECM organization and modification as well as cell adhesion compared to the S5-S7 MCT branch associated with ECM production, suggesting that tendon differentiation was more advanced than that of MCT (Figure 1E). A switch in biological activities was thus observed at the root of the lineage diversification, indicating that fibroblasts switched from a common Homeobox-based program to distinct ECM production programs.

Homeobox genes are associated with positional information during the establishment of the three limb axes (reviewed in Pineault and Wellik, 2014; McQueen and Towers, 2020), while extracellular matrix production is associated with CT fibroblast differentiation (reviewed in Nassari et al., 2017). To explore further this switch in biological activities, we analyzed the expression patterns of positional genes and lineage genes *in silico* and *in situ* (Figure 2). At E4, the positional genes (*LMX1B* for dorsal, *MSX2* for anterior, *HOXA11, HOXA13* for distal, *HOXD11* for posterior and *MEIS2* for proximal) showed regionalized and complementary expression patterns on the E4 UMAP (Figure 2A,E), indicative of a clustering of E4 limb cells according to positional cues. At the same stage, *SCX* and *OSR1* expression pattern did not show obvious regionalization in the E4 UMAP (Figure 2B), while these lineage genes were expressed in dorsal and ventral limb regions, as previously shown (Figure 2,C,D; Havis et al., 2016, Stricker et al., 2012). *SCX-* and *OSR1-*expressing cells therefore did not segregate according to their lineage cues but rather to limb positional information cues (Figure 2B-D). After E4, when the dorso-ventral and antero-posterior axes are established, the proximo-distal axis is still under construction as limbs still grow along this axis. *MEIS2* and *HOXA13,* as proxies of proximal and distal cues respectively (Nelson et al., 1996, Mercader et al., 2000), showed regionalized and complementary expression patterns up to E7 (Figure 2E). Their regionalization and complementary expression were lost after E7 (Figure 2E). At E6, we observed the emergence of fibroblast populations with no proximo-distal cue (Figure 2E, see ellipses). These regions correspond to overlapping expression of *OSR1, SCX* and *TWIST2* on E6/E7 UMAPs (Figure 2F,G). In E6 limbs, at the onset of spatial organization of the fibro- muscular system, *SCX* and *OSR1* expression was partially overlapping in and around muscle masses (Figure 2H-J). After E7, at E9 and E10, *SCX, OSR1* and *TWIST2* showed regionalized expression patterns on the UMAPs (Figure 2F,G). The perfect complementary expression of *SCX* and *OSR1* on the E10 UMAP (Figure 2F) and in limbs (Orgeur et al., 2018) is consistent with the transcriptional repression of *SCX* expression by OSR1 in limb cell cultures (Stricker et al., 2012). We conclude that from E7, *i.e* the onset of fetal period, fibroblasts cluster according to their lineage identity and no longer to their positional information.

**Figure 2.**
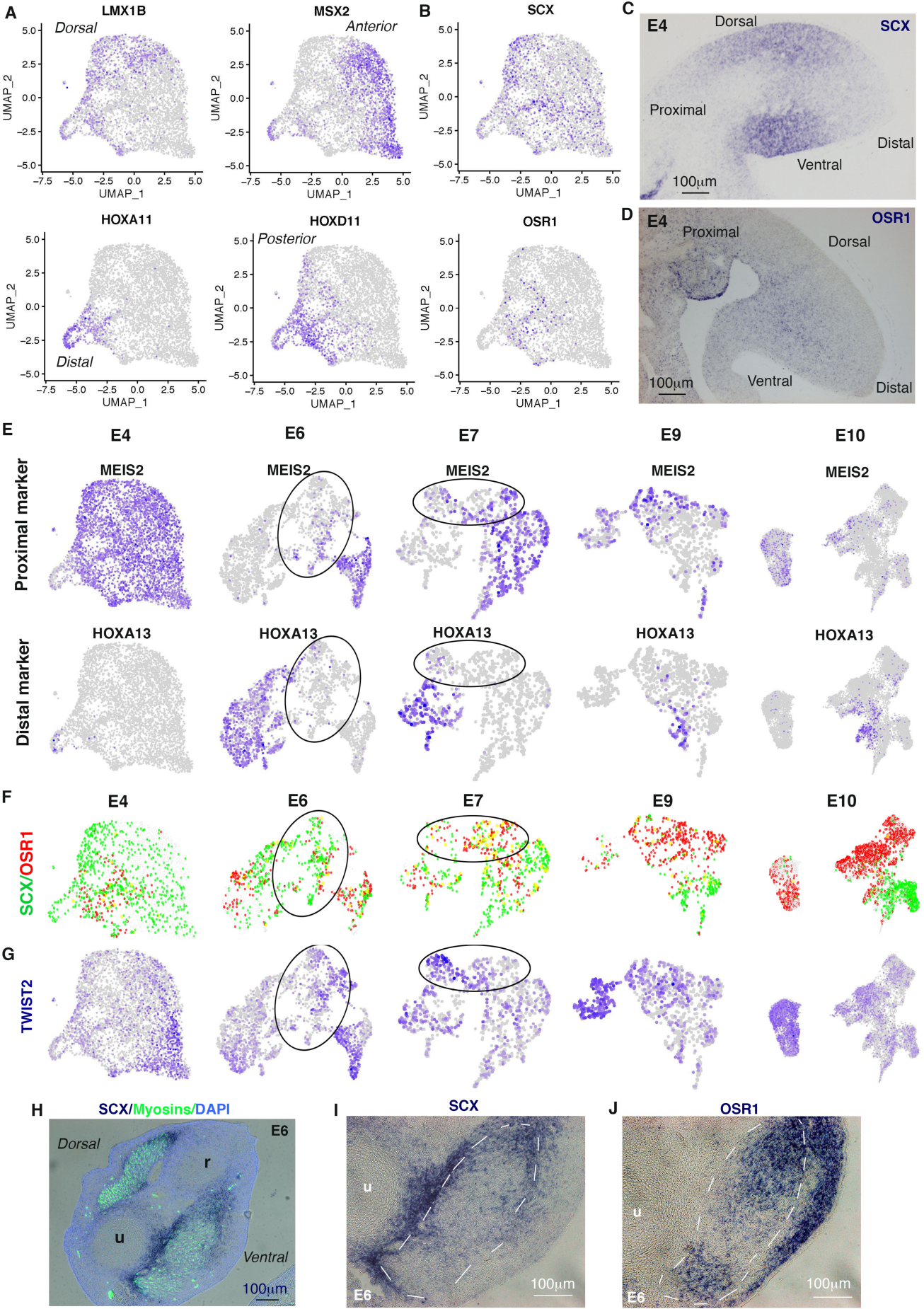
**CT fibroblasts switch from a positional information program to a lineage diversification program around E7**(**A**) Feature plots showing the distribution of cells positive for the positional markers *LMX1B, MSX2, HOXA11* and *HOXD11* in the E4 CT dataset. (**B**) Feature plots showing the distribution of cells positive for the lineage markers *SCX* and *OSR1* in the E4 CT dataset. (**C,D**) Colorimetric *in situ* hybridization with SCX (C) or OSR1 (D) probes to longitudinal limb sections of E4 embryos. (**E**) Feature plots showing the distribution of *MEIS2+* cells (proximal marker) and *HOXA13+* cells (distal marker) in E4, E6, E7, E9 and E10 CT datasets. (**F**) Feature plots showing the distribution of *SCX+* cells (green dots), *OSR1+* cells (red dots) and SCX+/OSR1+ cells (yellow dots) in E4, E6, E7, E9 and E10 CT datasets. (**G**) Feature plots showing the distribution of *TWIST2+* cells in E4, E6, E7, E9 and E10 CT datasets. Ellipses in (E, F, G) delineate UMAP regions with no positional information but emerging lineage information. (**H,I,J**) Colorimetric *in situ* hybridization with SCX (H,I) or OSR1 (J) probes (blue) to transverse limb sections of E6 chicken embryos. (H) Myosins and DAPI staining was overlayed over *SCX* blue labeling. (I,J) Ventral muscle masses are delineated with white dashed lines. u, ulna, r radius.

### Dermis, MCT and Tendon are composed of molecularly-distinct populations at E10

To identify potential fibroblast populations within the limb CT types, we focused on the E10 stage since the final fibro-muscular pattern is set at this stage. Unsupervised clustering analysis of the E10 CT dataset revealed nine distinct clusters (Figure 3A) and provided us with marker lists for each cluster, *i.e* lists of differentially expressed genes in one given cluster compared to the remaining clusters (Table S2). The analysis of *TWIST2, OSR1* and *SCX* expression on the E10 UMAP, combined with the localization of the nine E10 clusters on the 5CT trajectory, allowed us to attribute each cluster to a Dermis, MCT or Tendon identity (Figure 3A-C). We identified two dermal clusters, four MCT clusters and three tendon clusters. Dermis, MCT and tendon fibroblasts corresponded to 26.7%, 44.4% and 28.9% of all limb CT fibroblasts, respectively. The heatmap of the top 10 markers of each cluster, ranked by fold-change and ordered by CT type, showed that the clusters of Dermis and MCT types exhibited specific molecular signatures, while the signatures of Tendon clusters could be established but were more promiscuous (Figure 3D). We conclude that Dermis, MCT and Tendon types are subdivided in fibroblast populations with distinct but overlapping molecular signatures.

**Figure 3.**
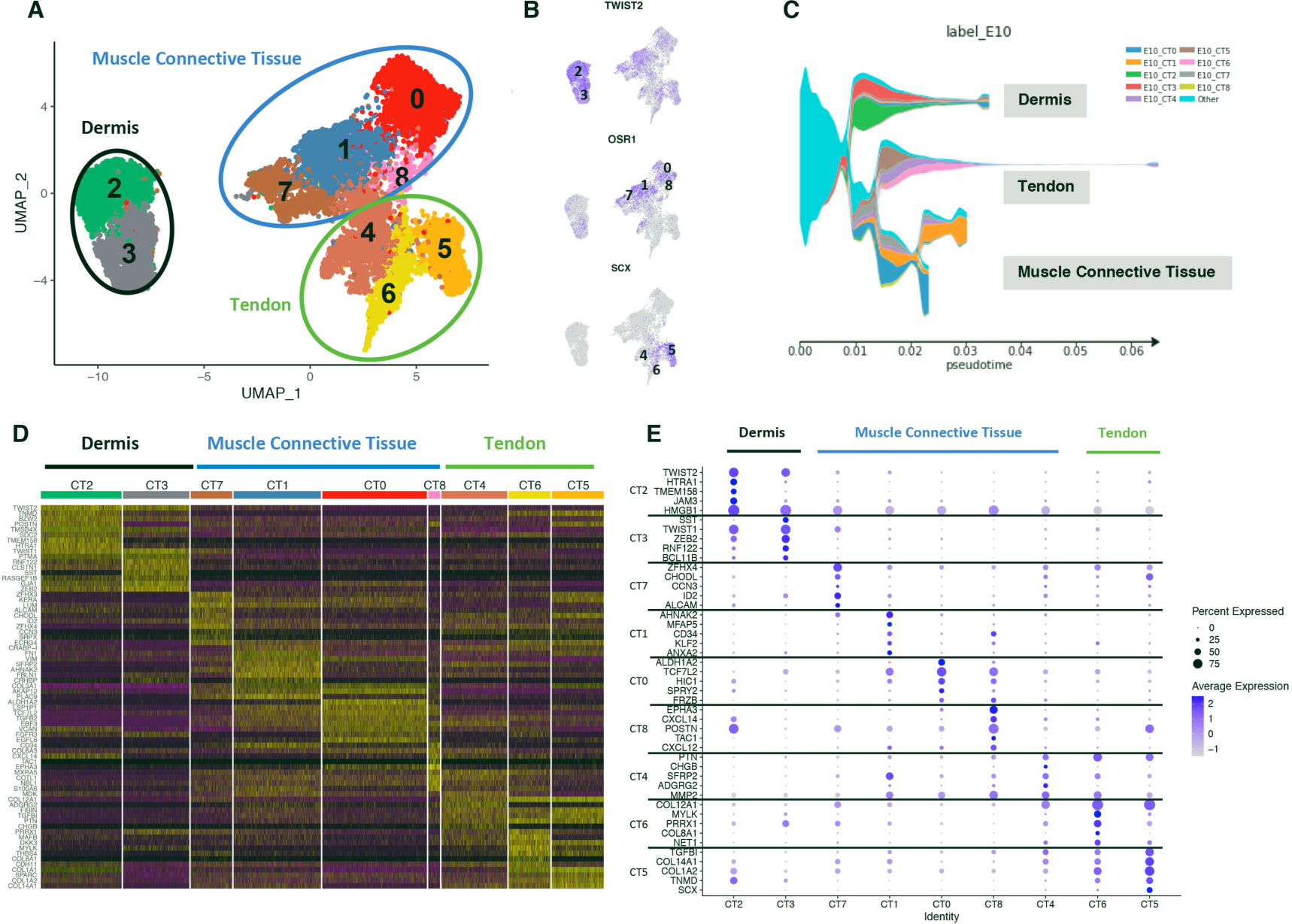
Dermis, MCT and Tendon are composed of molecularly-distinct populations at E10. (**A**) UMAP plot showing the distribution of the CT fibroblast clusters at E10. (**B**) Feature plots showing the distribution of *TWIST2+, OSR1+* and *SCX+* fibroblasts across CT clusters at E10. (**C**) Dendrogram of the 5CT inferred trajectory, highlighting the distribution of E10 clusters. Clusters from the other stages are grouped in light blue. (B) and (C) combined allow to position the Dermis, MCT and Tendon types on (A). (**D**) Heatmap showing the relative expression of the top 10 markers across all cells for each of the nine CT clusters, ordered by CT types at E10. Upregulated genes in yellow, downregulated genes in purple. (**E**) Dotplot showing the average expression levels and the fraction of expressing cells for a selection of genes in each of the 9 CT clusters, ordered by CT types.

In order to map the *in silico* clusters in fetal limbs, we selected up to four specific markers per cluster from the marker lists (Table S2) and characterized their expression patterns by *in situ* hybridization to E10 limb sections. Specific markers were chosen with the following criteria: association with a unique cluster, high fold-change value and high pct1 value (fraction of marker-expressing cells in the cluster of interest) combined to low pct2 value (fraction of marker-expressing cells in the other clusters) in order to select for the most specific markers. The dotplot shows a mix of top 5 markers, of single cluster-specific markers and of markers selected for ISH using the above criteria, ranked by fold-change and ordered by CT type, to illustrate the following general features (Figure 3E). Specific markers were expressed in at most 30% to 55% of the cells in the cluster of interest. Markers expressed in 90% to 100% of the cells for a given cluster, such as collagens, were non-specific markers, *i.e* they were expressed in a large fraction of cells in multiple clusters. For example, *COL1A1* and *COL3A1* were listed as markers for clusters 5-6 and 0-1-5, respectively (Table S2) and were also found in high proportions in other clusters (Figure S2A), consistent with collagen widespread expression in limbs (Lejard et al., 2011, Gaut and Duprez, 2016). In the following sections, we will describe each cluster, grouped by CT type, following the neighborhood order of the E10 UMAP and starting from the Dermis clusters.

### Dermal fibroblasts are divided in two molecularly-distinct populations with no obvious spatial regionalization

The Dermis type was divided into two transcriptionally-distinct populations (clusters 2 and 3), which was reminiscent of the two dermis layers, the upper papillary and lower reticular dermis (Lynch and Watt, 2018). *TWIST2*, a general marker for Dermis (clusters 2 and 3), was expressed in dermis of feather buds and inter buds, while being excluded from the ectoderm (Figure 4A-C). The *HTRA1* (HtrA Serine Peptidase 1) gene, a specific marker of cluster 2 (expressed in 52% of cluster 2 fibroblasts) showed a higher expression in inter-bud dermis than in feather bud dermis (Figure 4C,D,G,J). The *BCL11B* gene, a specific marker of cluster 3 (38% of cluster 3 fibroblasts), displayed enriched expression in dorsal dermis (Figure 4C,E,H,K). The *SST* (Somatostatin) gene, another specific marker of cluster 3 expressed in 42% of cluster 3 fibroblasts, was expressed in sub-regions of the feather bud dermis (Figure 4C,F,I,L). Analysis of additional cluster 2 markers, *TNMD* and *POSTN* (Figure 6L, Figures S6E), did not show any regionalization. Although the expression pattern of certain genes are highly regionalized, they did not allow us to assign clusters to specific regions such as the papillary and reticular dermis layers, or the inter bud and feather bud dermis, or the dorsal and ventral dermis. This is consistent with other scRNAseq studies that did not identify obvious molecular signatures for dermis regions (Lynch and Watt et al., 2018, Vorstandlechner et al., 2020).

**Figure 4.**
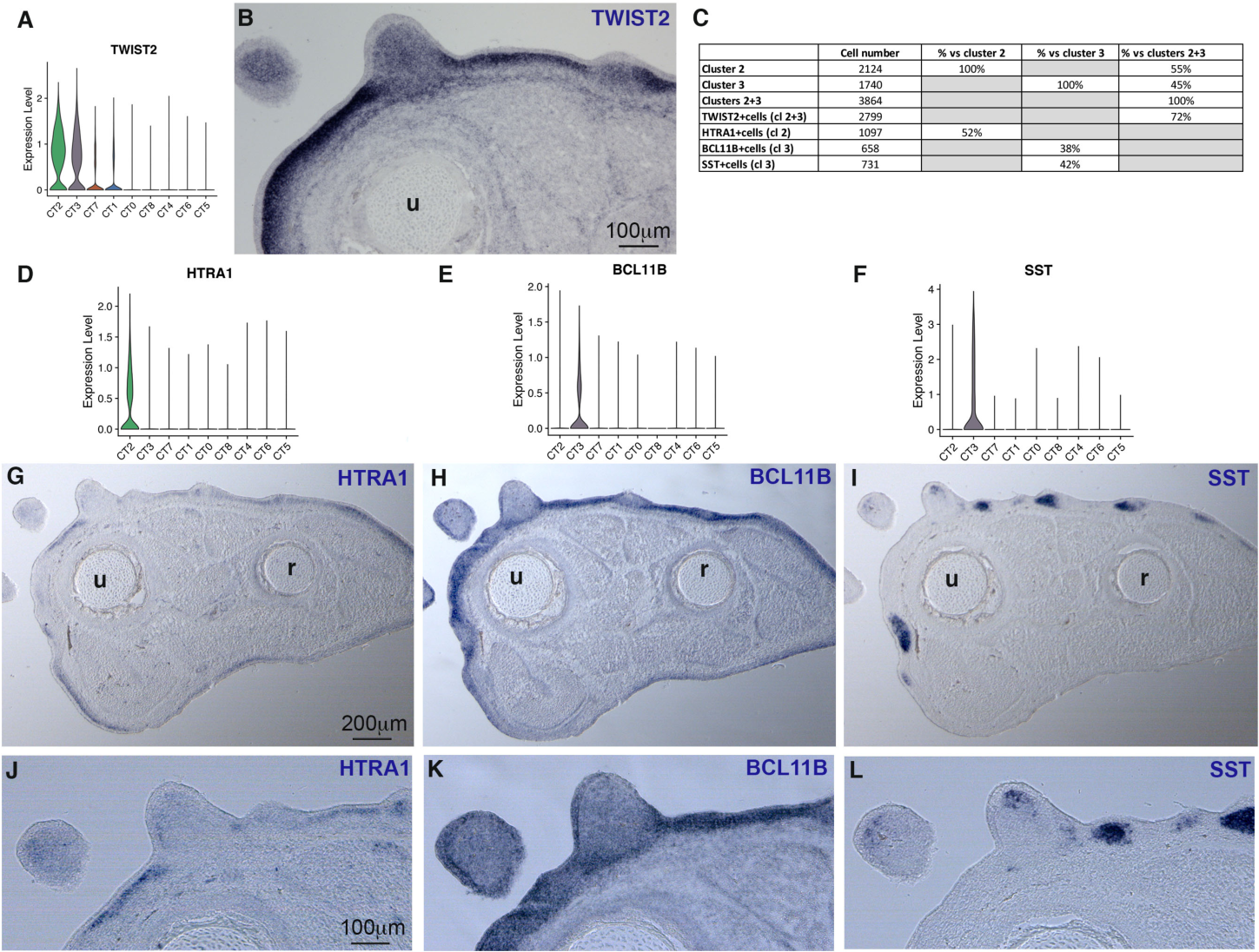
Dermal fibroblasts are divided in two molecularly-distinct populations with no obvious spatial regionalization. (**A,B**) *TWIST2* expression in CT clusters with violin plot of log-normalized expression levels in cells grouped by cluster (A) and in chicken limbs with *in situ* hybridization to transverse limb sections of E10 chicken embryos (B). (**C**) Cell numbers and associated percentage of dermal markers in clusters 2 and 3. (**D-F**) Feature and violin plots, respectively showing the distribution and log-normalized expression levels of *HTRA1* (D), *BCL11B* (E) and *SST* (F) genes. (**G-L**) Colorimetric *in situ* hybridization to adjacent limb transverse sections of E10 chicken embryos with HTRA1 (G,J), BCL11B (H,K) and SST (I,L) probes (blue labeling). (J,K,L) are high magnifications of dorsal and posterior dermis regions from sections shown in (G,H,I). Limb sections are oriented dorsal to the top and posterior to the left. u, ulna; r, radius.

### MCT clusters map to concentric fibroblast layers

The MCT type was divided into four fibroblast populations, clusters 7, 1, 0 and 8 (Figure 3A). The four MCT clusters corresponded to nearly half of limb CT fibroblasts (44.4%) and included the two largest fibroblast clusters (clusters 0 and 1). *PDGFRA*, *PRRX1* and *TCFL7L1* are recognized generic MCT markers routinely used to generate mouse Cre lines to study MCT (Yaseen et al., 2021, Sefton and Kardon, 2019). As expected, *PDGFRA, PRRX1* and *TCFL7L1* displayed a widespread expression both *in silico* and *in situ* with a noticeable *PRRX1* expression in tendon *in silico* populations and limb tendons (Figure S2B-J). Not only were those classical markers unsuitable to discriminate MCT fibroblast populations, but they were also not restricted to the MCT type in chicken fetal limbs.

*In situ* hybridization for selected markers allowed us to assign the four MCT clusters to distinct locations in limbs (Table S2, Figure 5). No obvious specific marker emerged for <u>cluster 7</u>. However, *CHODL* (Chondrolectin)*, ZFHX4* (Zinc Finger Homeobox 4) and *CCN3* (Cellular Communication Network 3) have in common to be markers of cluster 7, although being expressed in other clusters (Table S2). Their expression overlapped in a fibroblast layer underneath the dermis and surrounding the entire musculo-skeletal system (Figures 5A,E,I; S3), in a region that anatomically corresponds to the hypodermis. We conclude that cluster 7 fibroblasts correspond to the hypodermis.

**Figure 5.**
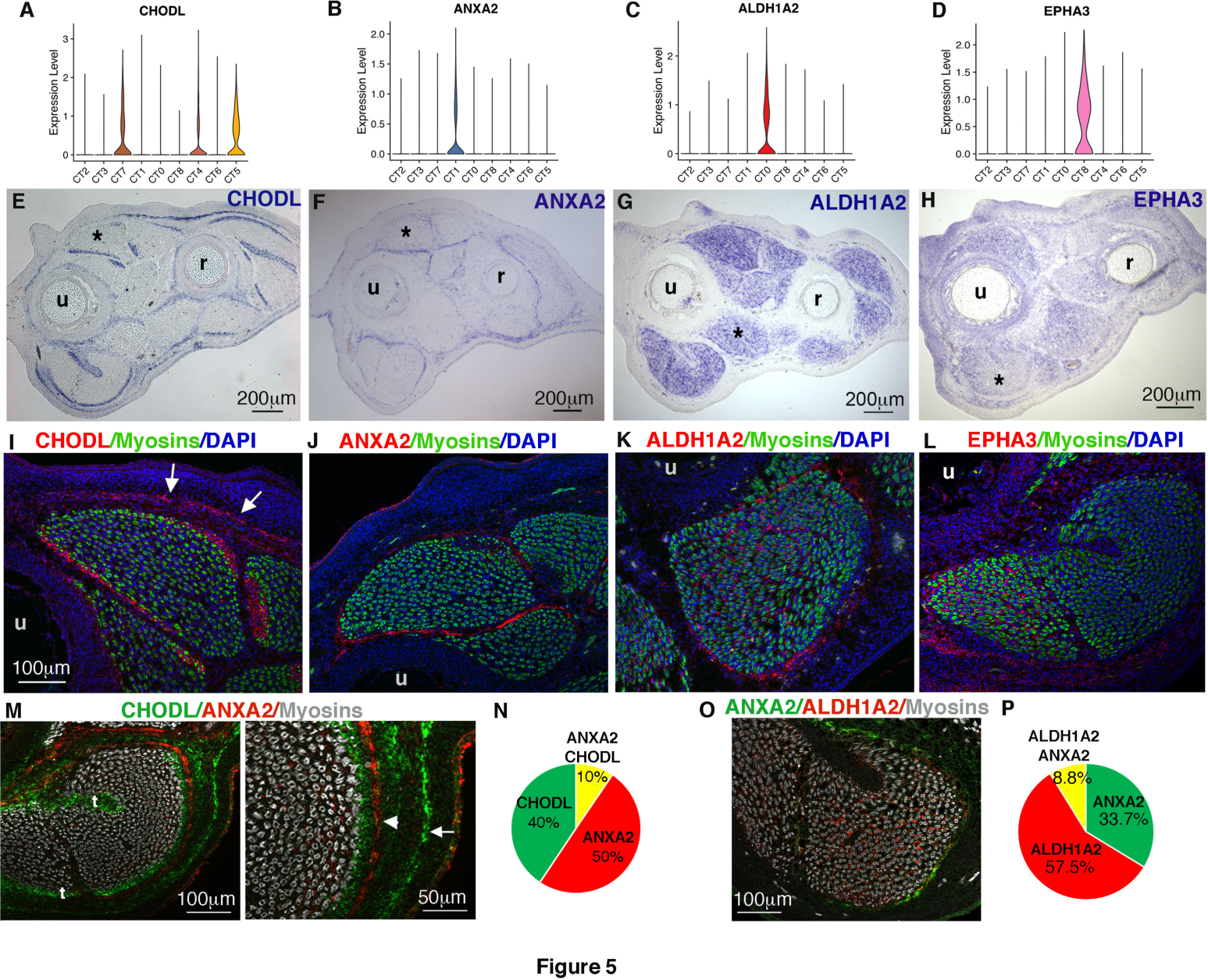
MCT populations display a hierarchical organization that prefigures the mature fibroblast layers in adult skeletal muscle. (**A-D**) Violin plots showing the log-normalized expression levels of one representative specific CT marker in cells grouped by cluster for each MCT cluster, *CHODL* for cluster 7 (A), *ANXA2* for cluster 1 (B), *ALDH1A2* for cluster 0 (C) and *EPHA3* for cluster 8 (D). (**E-H**) Colorimetric *in situ* hybridization to transverse limb sections at the level of the middle of the forearm of E10 chicken embryos with CHODL (E), ANXA2 (F), ALDH1A2 (G) and EPHA3 (H) probes (blue labeling). (**I-L**) Fluorescent *in situ* hybridization to transverse limb sections hybridized with CHODL (I), ANXA2 (J), ALDH1A2 (K), EPHA3 (L) probes (red fluorescent labeling) combined with immunostaining with the MF20 antibody recognizing myosins (green labeling). Limb muscles shown in (I,J,K,L) are labeled with black asterisks in limb sections of (E,F,G,H), respectively. (I) Arrows point to *CHODL* gene expression in the hypodermis. (**M**) Double fluorescent *in situ* hybridization to transverse limb sections focused on the FCU muscle (posterior and ventral muscle) with CHODL in green (cluster 7) and ANXA2 in red (cluster 1) probes. The white arrow on high magnification points to the hypodermis (green CHODL labeling), while the arrowhead points to epimysium (red ANXA2 labeling). (**N**) Percentage of *CHODL+*, *ANXA2+* and *CHODL+/ANXA2+* fibroblasts among the *CHODL-ANXA2* population. (**O**) Double fluorescent *in situ* hybridization to transverse limb sections focused to the FCU muscle with ANXA2 in green (cluster 1) and ALDH1A2 in red (cluster 0) probes. (**P**) Percentage of *ANXA2+, ALDH1A2+* and *ALDH1A2+/ANXA2+* fibroblasts among the *ANXA2-ALDH1A2* population. Limb sections are oriented dorsal to the top and posterior to the left. t, tendon, u, ulna; r, radius.

**Figure 6.**
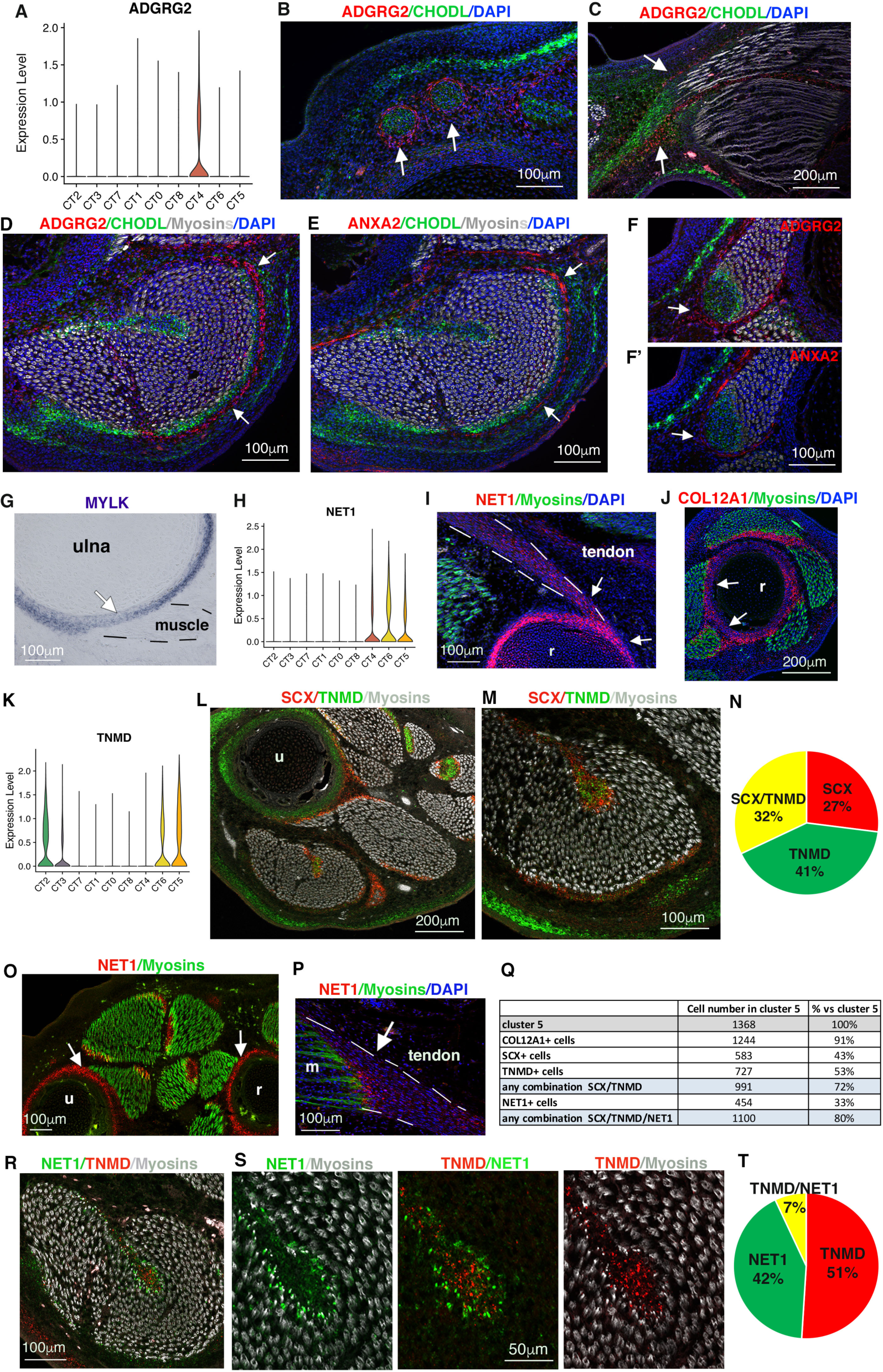
Peritenon, enthesis/perichondrium, and tendon proper correspond to distinct Tendon type populations (A) Violin plot showing the log-normalized expression levels of *ADGRG2,* a specific marker of tendon cluster 4, in cells grouped by clusters. (**B,C**) Double *in situ* hybridization with ADGRG2 probe (red) and CHODL probe (to label tendons in green) to transverse limb sections at distal level (B) and longitudinal limb sections (C) of E10 chicken embryos. (B,C) Arrows point to *ADGRG2* expression (red) surrounding tendons (green). (**D,E)** Double fluorescent *in situ* hybridization to adjacent transverse limb sections at E10 with ADGRG2 (red) /CHODL (green) (D) and ANXA2 (red) /CHODL (green) probes (E) followed with an immunohistochemistry with MF20 antibody recognizing myosins (greys) combined with DAPI staining. **(F,F’**) Double fluorescent *in situ* hybridization to adjacent transverse limb sections at E10 with ADGRG2 (red) /CHODL (green) (F) and ANXA2 (red) /CHODL (green) probes (F’) followed with an immunohistochemistry with MF20 antibody recognizing myosins (greys) combined with DAPI staining. (D-F’) White arrows point to *ADGRG2* (D,F) and *ANXA2* (E,F’) expression surrounding tendons and muscles. (**G**) Expression of *MYLK* gene, a marker of tendon cluster 6 with colorimetric *in situ* hybridization to transverse limb sections at E10 with *MYLK* probe (blue), focused on tendon attachment to cartilage element. (**H**) Violin plot showing the log-normalized expression levels of *NET1,* a marker of tendon cluster 6, in cells grouped by clusters. (**I**,**J**) Fluorescent *in situ* hybridization to longitudinal tendon section with NET1 probe (red) (I) and to transverse limb section with COL12A1 probe (red) (J) followed by an immunohistochemistry with the MF20 antibody that recognized myosins (green). (I,J) Arrows point to tendon attachments to cartilage elements labeled with *NET1* (I) and *COL12A1* (J). (**K**) Violin plot showing the log-normalized expression levels of *TNMD*, a marker of cluster 5, in cells grouped by clusters. (**L,M**) Double fluorescent *in situ* hybridization to transverse limb sections at E10 (L), focused on FCU muscle (M) with *SCX* (red) and *TNMD* (green) probes, followed with an immunostaining with the MF20 antibody (grey). (**N**) Percentage of *SCX+, TNMD+* and *SCX+/TNMD+* fibroblasts among the *SCX-TNMD* population. (**O,P**) Fluorescent *in situ* hybridization with the NET1 probe (red) followed with an immunostaining with the MF20 antibody (green) to transverse limb sections (O) and to longitudinal muscle sections (P) of E10 chicken embryos. Arrows point to *NET1* expression in tendons (O) and to tendon attachment close to muscle (P). (**Q**) Number of tendon fibroblasts in cluster 5. (**R,S**) Double fluorescent *in situ* hybridization to transverse limb sections at E10, focused on FCU muscle with NET1 (green) and TNMD (red) probes. (**T**) Percentage of *NET1+, TNMD+* and *NET1+/TNMD+* fibroblasts among the *NET1-TNMD* population.

*ANXA2* (the calcium regulated phospholipid-binding protein Annexin A2) was the main specific marker for <u>cluster 1</u> (Table S2). *ANXA2*, along with the additional specific markers, *AHNAK2, KLF2* and *CD34*, were expressed in a fibroblast layer delineating individual muscles and their associated tendons (Figures 5B,F,J; S4). Double *ANXA2/AHNAK2* fluorescent *in situ* hybridization confirmed that they were expressed within the same peripheral layer (Figure S4R,S). Cluster 1 fibroblasts delineate each individual limb muscle and thus prefigure the future epimysium.

The enzyme necessary for Retinoic Acid (RA) synthesis, *ALDH1A2*, was the main specific marker for <u>cluster 0</u> (Table S2). *ALDH1A2* transcripts labeled interstitial fibroblasts within limb muscles and were excluded from myosin+ cells (Figures 5G,K; S5). Three other specific markers for cluster 0 were expressed in the same location: *HIC1* (Hyper methylated in cancer1), a marker of quiescent fibroblasts in skeletal muscle (Scott et al., 2019) and the inhibitors of the FGF and Wnt signaling pathways, *SPRY2* and *FRZB* (Figure S5). We conclude that cluster 0 interstitial fibroblasts correspond to the future endomysium.

<u>Cluster 8</u> was the smallest fibroblast population and in close proximity to cluster 0 on the E10 UMAP (Figure 3A). Consistently, as for cluster 0, specific markers for cluster 8 were expressed in interstitial fibroblasts but enriched in a subset of limb muscles. The main specific marker *EPHA3*, along with the additional specific markers *POSTN* and *CXCL12,* were enriched in interstitial fibroblasts of the ANC muscle and of the posterior part of the FCU muscle (Table S2, Figures 5D,H,L; S6). The ventral part of the EMR muscle shows enriched *POSTN* and *CXCL12*, but not *EPHA4,* expression (Figure S6D,E,F). We could not correlate the cluster 8 molecular signature to slow and fast muscle type (Duprez et al., 1999).

To analyze the interface between MCT fibroblast populations, we performed double fluorescence *in situ* hybridization for *CHODL* (cluster 7) and *ANXA2* (Cluster 1) and for *ANXA2* (cluster 1) and *ALDH1A2* (cluster 0) and quantified the double-positive cells *in silico*. With both approaches, we find very little overlap between markers of distinct MCT clusters (Figure 5M-P), confirming that these clusters correspond to distinct fibroblast populations that are spatially segregated.

We conclude that the future hypodermis (cluster 7), epimysium (cluster 1) and endomysium (cluster 0) are molecularly distinct in E10 chicken limbs. The MCT populations display a concentric and hierarchical organization in fetal limbs that prefigures the mature fibroblast layers in adult skeletal muscle.

### Fetal tendons are divided in three fibroblast populations: peritenon, enthesis/perichondrium and tendon proper

Three clusters (clusters 4, 5 and 6) were assigned to Tendon identity (Figure 3A). Consistent with the promiscuity of their molecular signatures (Figure 3D), we identified markers common to the three tendon clusters in the marker lists, exampled with *COL12A1* (coding for the alpha1 chain of the FACIT type XII collagen), *PTN* (coding for the secreted molecule pleiotrophin) and *TGFBI* (coding for the secreted molecule, Transforming growth factor beta induced) (Table S2). *COL12A1, PTN* and *TGFBI* genes displayed similar expression domains associated with tendons as compared to *SCX* expression in limbs (Figure S7A-H). This is consistent with collagen XII role in tendon structure and function (Izu et al., 2021) and *PTN* expression in developing limb tendons (Mittapalli et al., 2009). Because the addition of all *COL12A1, PTN and TGFBI* gene combinations corresponded to 93% of cluster 4/5/6 fibroblasts (Figure S7H), we reasoned that their combined expression domains include the regions of the three tendon populations.

As for the MCT cluster 7, <u>cluster 4</u> displayed very few specific markers (Table S2). *ADGRG2* (adhesion G protein coupled receptor G2), a specific marker of cluster 4, was located in regions surrounding tendons (Figure 6A-F). *ADGRG2* was also observed delineating muscles in a similar manner to cluster 1 (epimysium) markers, exampled here with *ANXA2* (Figure 6D-F). Double fluorescent *in situ* hybridization against *ADGRG2* (cluster 4) and *ANXA2* (cluster 1) confirmed that both genes overlap in the fibroblast layer delineating muscle and associated tendons (Figure S7J,K). Double fluorescent *in situ* hybridization against *ADGRG2* (cluster 4) and *CHODL* (cluster 7, also expressed in tendons, Figure 5E) showed that *ADGRG2* was expressed underneath the cluster 7 hypodermis layer (Figure 6D). These results combined with the recent identification of *ADGRG* as a marker of horse peritenon (Pechanec and Mienaltowski, 2023) led us to conclude that cluster 4 fibroblasts correspond to the peritenon surrounding tendons and that peritenon and epimysium overlap when tendon interfaces with muscle.

Specific markers of <u>cluster 6</u> fibroblasts converged on gene expression in perichondrium and associated tendon attachments. *MYLK* (myosin light chain kinase), *COL8A1*, and *PRRX1,* involved in mouse perichondrium development (ten Berge et al., 1998), were strongly expressed in the perichondrium (Table S2, Figures 6G; S7L-Q; S2D,I). *NET1* (coding for neuroepithelial cell-transforming gene 1 protein) showed an expression in tendons close to the perichondrium similarly to *COL12A1,* a marker common to all tendon clusters (Figures 6H-J; S7A,D). We conclude that cluster 6 fibroblasts correspond to perichondrium and tendon attachments to perichondrium, namely the enthesis.

Specific markers for <u>cluster 5</u> pointed to recognized tendon genes involved in tendon development and homeostasis (Table S2). The top three markers of cluster 5 were known tendon-associated collagens, *COL1A1, COL12A1* and *COL14A1* (Lejard et al., 2011, Guerquin et al., 2013, Havis et al., 2014), albeit being listed as markers for other Tendon clusters, as well (Table S2). Other markers of cluster 5 known to be involved in tendon function included *POSTN* (coding for the secreted extracellular matrix protein, periostin) (Rolnick et al., 2022) and *SPARC* (coding for the secreted protein acidic and cysteine rich) (Wang et al., 2021). One striking observation was that the main recognized tendon markers, *SCX* and *TNMD,* were not expressed in the same tendon region; *TNMD* transcripts appeared to be enriched in the tendon core, while *SCX* transcripts were more homogenously distributed across the tendon (Figure 6K-N). In addition to being a specific marker of enthesis and perichondrium (cluster 6) (Table S2), *NET1* was expressed in cluster 5 tendon fibroblasts and in limb tendons (Figure 6O-Q). *NET1* expression was enriched in tendon fibroblasts in contact with or very close to muscle cells (Figure 6P). Double fluorescent *in situ* hybridization confirmed that *NET1+* cells were enriched in apposition to muscle cells and did not co- localize with *TNMD*+ cells that were more centrally located (Figure 6R,S), while *NET1+* cells overlapped with *SCX+* cells in this area (Figure S7O-Q). *In silico* quantification of double-positive cells further confirmed the little overlap between *NET1+* and *TNMD*+ cells (7% of *NET1+ TNMD+* population) (Figure 6T). We conclude that cluster 5 contains the fibroblasts of the tendon proper. Tendon proper fibroblasts are heterogeneous with fibroblasts located in the tendon core and others at the tendon periphery.

We conclude that the fetal tendon fibroblasts are divided into 3 populations, peritenon (cluster 4), enthesis/perichondrium (cluster 6) and tendon proper (cluster 5), which prefigure the mature tendon organization.

### CT fibroblast clusters are identified by a mosaic of partially overlapping gene expression patterns

Each fibroblast cluster is associated with a list of specific markers. However, as stated above, a specific marker for a given cluster is expressed in at most 30% to 55% of fibroblasts (Table S2). In addition, only 30% (on average) of the cluster fibroblasts are double-positive for any given couple of specific markers, while the addition of all the possible specific marker combinations accounts for 90% (on average) of the cluster cells (Figures 4C, 6Q, S3M,N, S4Q,R, S5Q,R, S6M,N, S7,H,I,P). This shows that a fibroblast cluster corresponds to a combination of genes with partially overlapping expression patterns. Therefore, these fibroblast populations cannot be defined by the expression of one key gene but by the combined expression of multiple genes.

### MCT and Tendon fibroblast populations successively emerge from a common population at the onset of the fetal period

In order to identify the developmental origin of the E10 CT clusters and their time of emergence, we performed trajectory inference analysis with STREAM (Chen et al., 2019) on datasets resulting from the integration or the merge of two consecutive CT datasets between E6 and E10 (Figures 7, S8). E4 was excluded from the analysis since it did not contain lineage-relevant information. The analysis of the inferred trajectory for each combined dataset enabled us to establish a lineage link between individual clusters of successive stages (Figure S8A-C), which when compiled, leads to a lineage tree of individual clusters across time (Figure S8D). To better visualize the lineage tree of MCT and Tendon clusters, we duplicated the lineage tree for the E9 and E10 stages (shown in Figure S8D) and highlighted the MCT- and Tendon-related trajectories (Figure 7).

**Figure 7.**
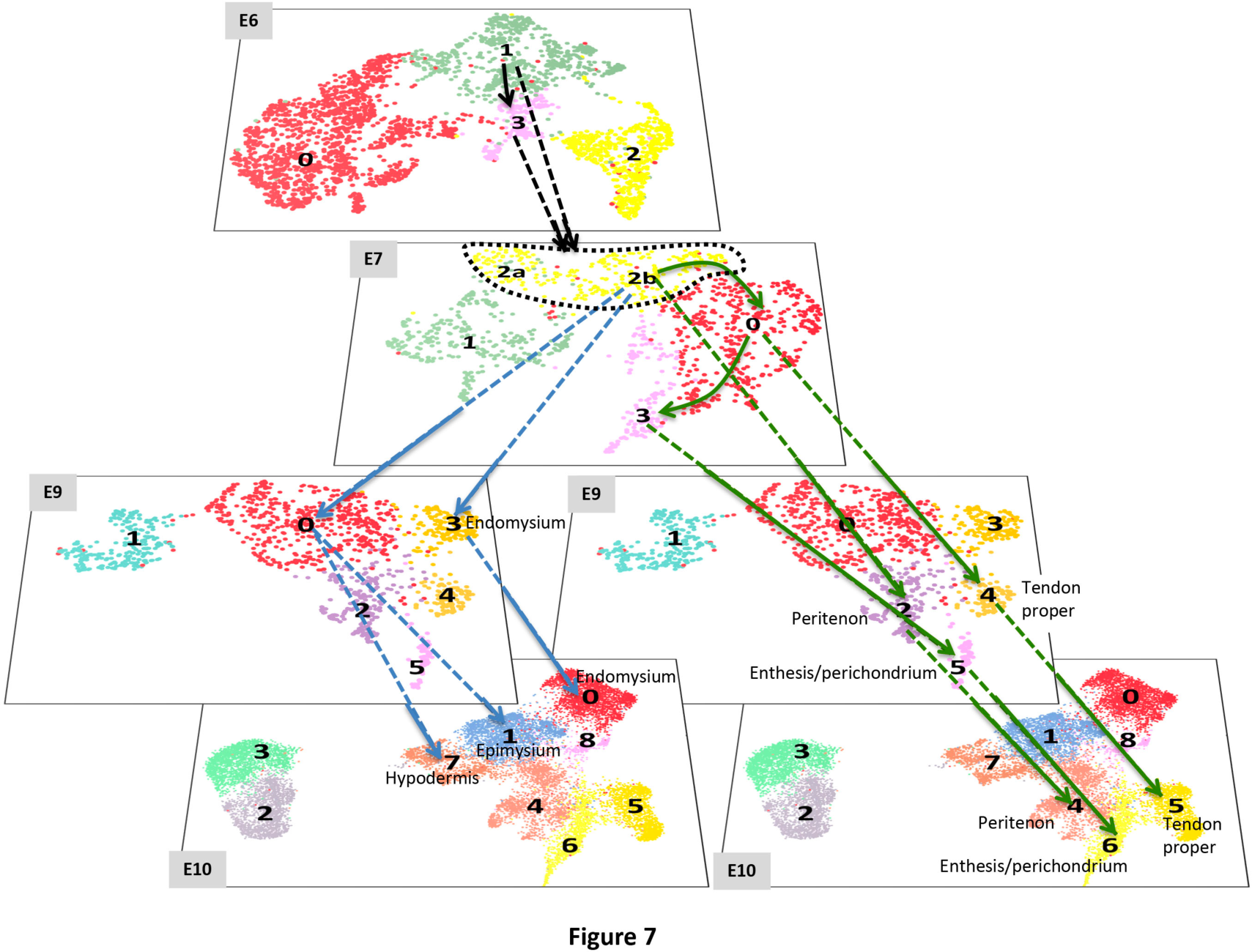
Lineage tree of CT clusters focused on the MCT and Tendon lineages. Representation of the CT cluster lineage tree derived from the analysis of STREAM trajectories performed on the E6-E7, E7-E9 and E9-E10 combined CT datasets. UMAP plots showing the distribution of CT clusters for the E6, E7, E9 and E10 CT datasets. To illustrate that Tendon and MCT lineages segregate from E7, the cluster lineage tree was separated in two branches and the E9 and E10 UMAP plots were duplicated. Black arrows refer to trajectories before lineage segregation. Blue arrows refer to MCT-related trajectories. Green arrows refer to Tendon-related trajectories. Trajectories unrelated to the MCT and Tendon lineages, such as Dermis-related trajectories, are not shown. The cluster identity is indicated when identified.

At E7, consistently with the 5CT trajectory (Figure 1B), the Dermis type clusters (E10- clusters 2, 3) were among the first fibroblast populations to emerge. We identified its origin in a sub-region of E7-cluster 2 labeled cluster2a. The E7-cluster 2b was identified as the common origin for the MCT and Tendon types. Consistently, E7 was the stage where the proportion of *OSR1*+/*SCX*+ double-positive cells was the highest and the highest proportion of those *OSR1*+/*SCX*+ double-positive cells was found in E7-cluster 2b (Figure S9A,B). These results support the existence of a common progenitor for both CT types in limbs, which is consistent with *Scx*- or *OSR1*-derived fibroblasts contributing to MCT or tendon, respectively (Esteves de lima et al., 2021). At E9, the endomysium (E10-cluster 0) was the first MCT cluster to emerge from E7-cluster 2b. This is consistent with the splitting of the MCT branch at E9 on the 5CT trajectory (Figure 3C). At E10, the last MCT clusters to emerge were the epimysium (E10-cluster 1) and hypodermis (E10-cluster 7), both deriving from the splitting of E9-cluster 0. Concomitantly to MCT cluster emergence, the enthesis/perichondrium (E10-cluster 6) was the first Tendon cluster to emerge as E7-cluster 3 deriving from E7-cluster 0. By E9, the other two Tendon clusters (E10-clusters 4, 5) have emerged from the E7-cluster 2b and the E7-cluster 0, respectively.

We conclude that the MCT and Tendon fibroblast populations successively emerge from a common population at the onset of the fetal period.

## Discussion

In this work, we showed that limb fibroblasts successively differentiate into CT populations after loosing their positional information at the onset of fetal period. When the final fibro- muscular pattern is set, transcriptionally-distinct but overlapping fibroblast populations are regionalised in limbs, prefiguring the mature fibroblast layers associated with adult skeletal muscle.

### Limb fibroblasts provide positional cues and differentiate in a sequential manner

Fibroblasts provide the positional information cues to the limb cells during the establishment of the three limb axes (Pineault and Wellik, 2014; McQueen and Towers, 2020). This is clear from our gene profiling analysis as fibroblasts cluster according to positional cues up to E7. At E7, fibroblasts switch to a differentiation program to CT types, indicating that they fulfil their two roles (provider of positional cues and differentiation) in a sequential manner. After this switch, they seem to lose their positional cues. At least, this positional program is no longer dominant in their transcriptional identity. However, we do observe residual *HOX* expression at E10 with *HOXA13* expression in cluster 4/peritenon (Figure 2D). Peritenon fibroblasts are known to display stemness features as compared to tendon proper fibroblasts and are required for tendon repair after injury (Dyment et al., 2013; Mienaltowski et al., 2013). *HOXA13* expression in peritenon fibroblasts could be associated with residual positional information maintained in immature fibroblasts only (Pineault and Wellik, 2014) or with the differentiation of the peritenon lineage; these possibilities are not mutually exclusive.

### CT fibroblasts differentiate without continuous recruitment of progenitors

As stated above, the second phase of limb CT development corresponds to differentiation and lineage diversification. A striking observation was that at each stage, CT clusters correspond to different populations, each at a given differentiation step. At each developmental stage, we did not identify clusters corresponding to successive differentiation steps. This is different from the muscle differentiation process, where at each developmental stage the different steps of myogenesis from progenitors to differentiated cells are observed as clusters in scRNAseq analysis of limb muscle cells (Esteves de Lima et al., 2022). For CT, one has to combine successive developmental stages to visualize the progression of CT fibroblast populations along the differentiation path. CT fibroblast clustering therefore highlights lineage diversification rather than fibrogenesis progression. This shows that muscle and CT form and grow in two different ways. Muscle grows by persistent recruitment of progenitors into the myogenic program, while CT progresses from one differentiation step to the next in cohorts, exhausting populations from the previous differentiation steps. CT growth may occur with cell proliferation for each fibroblast population at different differentiation steps. Indeed, the distribution of cell cycle phases shows a persistent mix of cell cycle phases up to E10. CT fibroblasts start to differentiate and diversify at E7, the transition stage from embryonic to fetal development when muscle contraction starts in chicken embryos. Interestingly, Coren and colleagues (2024, back-to-back submission) show in mouse limbs that fibroblast diversity arises at the same embryonic to fetal transition stage and depends on muscle contraction.

### Delineating and interstitial fibroblasts display distinct transcriptional identities in line with their functions

At fetal stages, chicken limb MCT contains two main fibroblast populations, delineating and interstitial fibroblasts that prefigure the future epimysium and endomysium of adult skeletal muscle (Figure 8). Strikingly, similar delineating and interstitial fibroblasts have been identified in developing and postnatal mouse limb muscles (Coren et al., 2024, back-to-back submission). Furthermore, inferred trajectories show that these two MCT populations derive from a common origin both in chicken (our study) and mouse (Coren et al., 2024). We further show in chicken limbs that interstitial fibroblasts (endomysium) emerge before delineating fibroblasts (epimysium). Nonetheless, once established, the epimysium contributes to the endomysium, *i.e* the peripheral layer feeding the more central layer (Figure 7, Figure S8). We believe that the perimysium (surrounding muscle bundle) is not present at fetal stages. Whether perimysium originates from epimysium or endomysium is not known.

**Figure 8.**
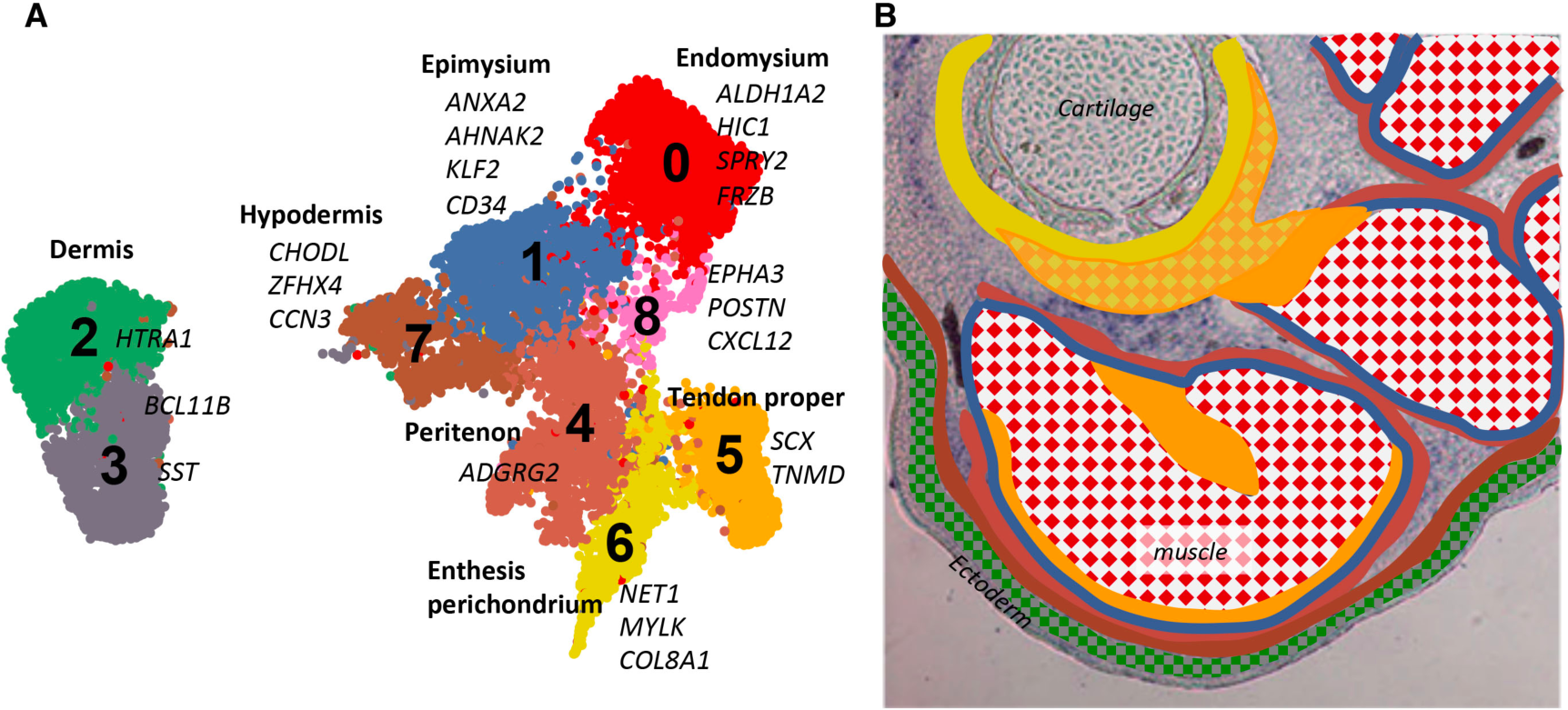
Spatial arrangement of fibroblast populations prefigures the mature fibroblast layers associated with adult skeletal muscle. (**A**) UMAP plot showing the distribution of CT clusters at E10. Specific markers for which *in situ* hybridization was performed are assigned to clusters. (**B**) Schematic summarizing the location of fibroblast populations on limb transverse section at E10. The same color code was used for the *in silico* clusters and the fibroblast populations in limbs, except for cluster 8, color-coded as cluster 0 for the sake of clarity. Dermal fibroblast populations (clusters 2,3) were located at the limb periphery underneath the ectoderm. Underneath the dermis, the hypodermis (cluster 7) a fibroblast layer that surrounds the whole musculoskeletal system. The fibroblast layers delineating tendon and muscles are the peritenon (Cluster 4) and epimysium (cluster 1) in continuity. In addition to the peritenon fibroblasts, tendon fibroblasts were divided into two transcriptionally-distinct clusters that correspond to tendon attachments to cartilage including perichondrium (cluster 6) and tendon proper (cluster 5). The interstitial fibroblasts (cluster 0 and 8) correspond to the future endomysium.

The function of the outermost CT sheath of muscle is to constrain muscle in size and shape and to prevent friction between muscles during contraction (Purslow, 2020). One main marker of these fetal delineating/epimysial fibroblasts is *ANXA2*. ANXA2 has been shown to be associated with membrane repair, cell-to-cell contact, membrane ruffling and cytoskeletal scaffolding (Lim and Hajjar, 2021; Méndez-Barbero et al., 2022), consistent with a function in maintaining/constraining muscle size. Moreover, ANXA2 partner proteins, such as the scaffold protein AHNAK2 (Pascal et al., 2022) and the Ca^2+^-binding protein S100A6 (Cheng et al., 2005), are also markers of delineating fibroblasts. ANXA2 promotes FAP differentiation towards adipogenesis in the context of limb girdle dystrophy (Hogarth et al., 2019). CD34, a marker of delineating fibroblasts, is also a marker of adipose stem cells (Raajendiran et al., 2019). Altogether, this is consistent with the epimysium as being a reservoir of adipocyte progenitors (Giulani et al., 2021). Interestingly, we found that delineating fibroblast population (cluster 1) has a common origin with hypodermis fibroblast population (cluster 7) that is also recognized to contain white adipose tissue progenitors (Lynch and Watt et al., 2018).

The interstitial/endomysial fibroblasts (cluster 0) found in between muscle fibers, have a Retinoic Acid (RA) signature, with *ALDH1A2* as the main marker of these fibroblasts. This enzyme is required for RA function, suggestive of RA activity in fetal interstitial fibroblasts. RA has been associated with FAP differentiation towards fibroblasts (and not adipocytes), while loss of RA signalling drives FAPs towards adipogenesis (Zhao et al., 2020). RA regulates matrix production and is seen as a therapeutic option to fight against fibrosis in skin disorders and to promote organ repair (reviewed in Wang et al., 2020). HIC1, another marker of interstitial fibroblasts, is a downstream target of RA (Kristensen et al., 2012) and has been shown to maintain muscle-associated fibroblasts in quiescence in adult normal and regenerative muscles (Scott et al., 2019). Inhibitors of FGF (Sprouty2) and Wnt (FRZB) signalling are also markers of interstitial fibroblasts. Interestingly, as for RA, Wnt inhibitors are also considered as a therapeutic option against fibrosis (Burgy et Königshoff 2018). We therefore see the hypodermis and epimysium external layers as a source of adipogenic progenitors, while the endomysium contains fibrogenic progenitors.

### Fetal tendon fibroblasts at different maturation states are spatially segregated

Fetal tendon contains three main fibroblast populations, the peritenon (cluster 4), enthesis/perichondrium (cluster 6) and tendon proper (cluster 5) that prefigure adult tendon organization (Figure 8). Enthesis/perichondrium (cluster 4) is the first population to emerge at E7, consistent with enthesis function that is recognized to drive tendon position along cartilage elements (Blitz et al., 2009, 2013). Enthesis/perichondrium fibroblast population is associated with adhesion molecules, exemplified by the adhesion molecule NET1. NET1 participates in cardiac fibrosis by promoting collagen synthesis in fibroblasts via the activation of Wnt/β-catenin and TGF/Smads signaling pathways (Li et al., 2023). Within the tendon proper (cluster 5), we identified a regionalization of tendon fibroblasts with the classical tendon markers *SCX* and *TNMD*. Because *TNMD* is associated with differentiated tendon fibroblasts while *SCX* expression is associated with immature tendon fibroblasts (Shukunami et al., 2006, 2018), we assume that tendon fibroblasts located in the tendon core are more advanced in differentiation than those in the tendon periphery. Peritenon (cluster 4), the outermost tendon layer, is seen as a type of synovial sheath allowing tendon gliding and is recognized to concentrate blood vessels and innervation necessary for tendon function (Purslow, 2020), consistent with the specific marker *GHGB* (chromogranin B), a secreted peptide of the neuroendocrine system (Helle et al., 2010). Tendon progenitors recruited for tendon repair are concentrated in the peritenon versus tendon proper (Dyment et al., 2013; Mienaltowski et al., 2013), suggestive of an immature state of this fibroblast layer. Consistently, at E10, peritenon (cluster 4) provides cells to the other two tendon clusters 6 and 5 (Figure S9C).

### A continuum in fibroblast identities to bridge tissues of highly different nature

Strikingly, each *in silico* fibroblast cluster maps to a specific *in situ* location in limbs, which supports the biological relevance of this scRNAseq approach (Figure 8). As stated in the Results, a fibroblast cluster corresponds to a mosaic of partially overlapping gene expression patterns. In addition, a gene identified as a marker for a given cluster by the differential gene expression analysis often shows weaker expression, outside of this cluster, both *in silico* and *in situ*. Along the same line, except for the dermis clusters that are more distant, CT clusters are in contact with at least two neighboring clusters, highlighting their transcriptional proximity. Finally, the neighbourhood relationships are conserved between the *in silico* and *in situ* analyses. Clusters that are neighbours on the E10 UMAP correspond to populations that are neighbours in limbs. This means that fibroblast populations that are close in transcriptional identity are also close in space. This shows that at the limb level, there is a continuum of promiscuous fibroblast identities reflecting the interconnected 3D network of fibroblasts associated with muscles. Consistently, our results point to a continuity between the outermost layers of tendon (peritenon) and MCT (epimysium). We propose that this continuum in identities underlies a continuum in mechanical properties that is instrumental to CT function. CT bridges muscle to bone or muscle to skin that are mechanically different. Our hypothesis is that it would do so by apposing CT layers with mechanical properties that would be close to that of muscle and progressively evolve towards that of bone, for example. The characterization of mechanical properties of fibroblast populations awaits further studies.

## MATERIALS AND METHODS

### Chicken embryos

Fertilized chicken (Gallus gallus) eggs from commercial sources (JA 57 strain: Morizeau, Dangers, France) were incubated at 38.5°C in a humidified incubator until appropriate stages. Embryos were staged according to the number of days *in ovo* (E). All experiments on chicken embryos were performed before E14 and consequently are not submitted to a licensing committee, in accordance with European guidelines and regulations.

### Seurat clustering analysis

#### Sample preparation

The sample collection and scRNAseq protocol from E7 and E9 limb cells were performed as described for E4, E6 and E10 by Esteves de Lima et al. (2021). Briefly, scRNAseq datasets were generated from whole forelimbs from one E7 embryo and one E9 embryo using the 10X Chromium Chip (10X Genomics) followed by sequencing with a HighOutput flowcel using an Illumina Nextseq 500 and by sequence analysis with Cell Ranger Single Cell Software Suite 3.0.2 (10X Genomics). Ectoderm at all stages and digits at E10 were manually removed before cell dissociation. Cartilage tissues were poorly dissociated and therefore excluded from the samples by the single-cell isolation protocol.

#### Seurat clustering of whole limb datasets

The Seurat package (v3.2.3) (Stuart et al., 2019) under R (v4.3.1) (R Core Team, 2019) was used to perform downstream clustering analysis on scRNAseq data (Macosko et al., 2015) (see Table S3 for parameter and cut-off values). Cells went through a classical Quality Control using the number of detected genes per cell (nFeatures), the number of mRNA molecules per cell (nCounts) and the percentage of expression of mitochondrial genes (pMito) as cut-offs. Outliers on a nFeature vs nCount plot were manually identified and removed from the dataset. Potential doublets were identified by running the Scrublet algorithm (Wolock et al. 2019) and then removed from the dataset. Gene counts for cells that passed the above selections were normalized to the total expression and Log-transformed with the NormalizeData function of Seurat using the nCount median as scale factor. Highly variable genes were detected with the FindVariableFeatures function (default parameters). Using highly variable genes as input, principal component analysis was performed on the scaled data in order to reduce dimensionality. Cell cycle effect was regressed out using the ScaleData function. Statistically significant principal components were determined by using the JackStrawPlot and the ElbowPlot functions. Cell clusters were generated with the FindNeighbors/FindClusters functions (default parameters except for the number of selected PCs). Different clustering results were generated at different resolutions and for different sets of PCs. Non-linear dimensional reduction (UMAP) and clustering trees using Clustree (Zappia et al. 2018) were used to visualize clustering results and select for the most robust and relevant result. Markers for each cluster were found using the FindAllMarkers function of Seurat (using highly variable genes as an input, default parameters otherwise) that ran Wilcoxon rank sum tests (p-val adjusted < 0.05). The clusters identified as CT clusters by the differential expression of classical CT markers (SCX, OSR1, TWIST2) represented the majority of limb cells, in addition to other clusters encompassing the expected cell populations present in developing limb tissues such as muscle, vessels/blood, neural crest cells and ectoderm (Figure S1A,B and Esteves de Lima et al., 2021).

#### Seurat clustering of CT subsets

For all datasets, we selected the CT cluster cells and extracted them from the post-Quality Control object to generate a CT subset of the whole limb dataset. CT datasets were then processed again through the pipeline described above, from the Normalization to clustering and marker identification steps. The CT datasets were analyzed using classical Seurat tools such as Feature plots, Violin plots, Dimplots, Heatmaps and Dotplots. Gene expression was defined by ‘gene log-normalized count>0’. The R package ggplot2 v3.4.4 (Wickham, 2016) was used to generate custom feature plots highlighting gene co-expression. Population intersection plots were generated with the R package UpSetR v1.4.0 (Conway et al., 2017).

#### Merge and RPCA integration of CT subsets

In order to conduct trajectory inference analyses, individual post-clustering CT datasets were combined into single datasets either using the Seurat Merge tool (E7-E9 merge) or the RPCA integration strategy after merging (5CT, E6-E7, E9-10 integrations) to deal with the batch effect observed between the E4/E6/E10 and the E7/E9 batches. Before RPCA integration, merged CT datasets were individually normalized, highly variable genes identified and used to select integration features with the SelectIntegrationFeatures function (default parameters). Scaling followed by PCA with cell cycle gene regression were performed on the integration features. Anchor cells were identified with the FindIntegrationAnchors function, using the integration features and the RPCA reduction (default parameters). CT datasets were integrated with the IntegrateData function, using the integration features and anchor cells (default parameters). The merged or RPCA-integrated objects were then processed through the pipeline described above, from the Normalization (merge) or the Scaling (RPCA) steps to the non-linear dimension reduction (UMAP) step before being imported into the STREAM environment.

### STREAM trajectory inference on merged/integrated CT datasets

The STREAM pipeline (v1.0) (Chen et al. 2019) under Python (v3.7.9) was used to perform trajectory inference analysis on integrated/merged CT datasets (see Table S3 for parameter values). The merged E7-E9 and the integrated E6-E7 and E9-E10 datasets were imported into the STREAM environment under the h5ad format to build the trajectory on the Seurat UMAP dimension reduction. To perform differential gene expression analysis on top of trajectory inference, the integrated 5CT dataset was imported under the 10X format and we generated an anndata object combining the unmodified expression matrix (RNA slot) of the 5CT Seurat object with a dimension reduction matrix derived from the integrated expression matrix of the 5CT Seurat object. The dimension reduction matrix was calculated on the integration features with the dimension_reduction function (method = ’mlle’, n_neighbor = 250). Trajectories were inferred with the elastic_principal_graph function, using the mlle reduction for 5CT and the umap reduction for the other datasets. Trajectories were visualized as a dendrogram, color-coded with the Seurat cluster labels or gene expression levels, with the plot_stream function. Variable genes and then branch-associated marker (z-score > 1.0; p value < 0.01) were identified from the unmodified expression matrix by the select_variable_genes and detect_leaf_markers functions (default parameters) for the 5CT.

### DAVID analysis of the branch-associated markers for the 5CT trajectory

GO analysis of the branch-associated markers was performed using the Functional Annotation Clustering tool from DAVID (Database for Annotation, Visualization and Integrated Discovery, version 2021, DAVID Knowlegebase v2023q4) (Sherman et al., 2022, Huang et al., 2009). GO analysis was performed on marker lists associated with the Dermis, MCT, SCX and origin branches. Selected categories (enrichment score > 2.0 ; p value < 0.05) were associated with GO terms grouped as follows: DNA binding, Homeobox and Transcription regulation ; Secreted, Glycoproteins and Extra Cellular Matrix ; Collagens, Extra Cellular Matrix organization and Fibrillar collagen ; Cell adhesion, Focal adhesion and Proteoglycans.

### *In situ* hybridization

Forelimbs of E6 or E10 chicken embryos were fixed in 4% paraformaldehyde (Sigma-Aldrich) overnight at 4°C, then processed in 7.5%/15% gelatin/sucrose (Sigma-Aldrich) for situ hybridization to cryostat sections, as previously described (Esteves de Lima et al., 2016, 2021, 2022). Transverse or longitudinal limb sections were performed. Alternating serial sections were hybridized with probe 1, probe 2, probe 3 and probe 4 to allow comparison of expression domains on adjacent sections of the same limb. Digoxigenin-labelled or Fluorescein-labelled mRNA probes were either prepared as PCR probes with primers listed in Table S4 using the Kit GoTaq green master mix (Promega) or as described (Orgeur et al., 2018, Esteves de Lima et al., 2021, 2022) for SCX, OSR1, KLF2, SPRY2 probes. Colorimetric *in situ* hybridization was performed according to Esteves de Lima et al. (2021, 2022) using an antibody against Digoxigenin labelled with alkaline phosphatase. Fluorescence *in situ* hybridization was performed according to Wilmerding et al. (2021), using antibodies (against Digoxigenin or Fluorescein) labelled with POD. Fluorescence was revealed with the TSA+ Cyanin 3 or 5 system (AKOYA biosciences).

### Immunohistochemistry

*In situ* hybridization was followed with an immunohistochemistry with the MF20 monoclonal antibody (DSHB cat. # MF20, undiluted supernatant), which recognized sarcomeric myosin heavy chains and allow the visualisation of differentiated muscle cells. The monoclonal antibody MF20, developed by A. Kawakami, was obtained from the Developmental Studies Hybridoma Bank developed under the auspices of the NICHD and maintained by the University of Iowa, Department of Biology Iowa City, IA 52242, USA. Secondary antibodies were conjugated with Alexa 488, Alexa 555 or Alexa 647 (Invitrogen). Nuclei were visualized with DAPI (Sigma-Aldrich) staining.

### Image capture

After immunohistochemistry or *in situ* hybridization experiments, images were obtained using a Zeiss apotome epifluorescence microscope or a Nikon eclipse E800 microscope, with the possibility to combine colorimetric and fluorescence labelling, or a leica DMR microscope for colorimetric images only.

## Acknowledgements

We thank Marie-Claire Delfini for critical reading the manuscript. This work was supported by the CNRS, Inserm, SU, ANR_LimbCT, ANR_MyoDom and AFM_MyoReg. ARTbio was supported by the CNRS, SU, the Institut Français de Bioinformatique (IFB) and by a grant from the SIRIC CURAMUS.

## Author contributions

Conceptualization: E.H., D.D.; Methodology: E.H., C.B., M.-A.B., L.B., D.D.; Software: E.H., L.B., T.G.; Validation: E.H., C.B., M.-A.B., D.D.; Formal analysis: E.H., D.D.; Investigation: E.H., C.B., D.D.; Resources: E.H., C.B., M.- A.B., D.D.; Data curation: E. H., D.D.; Writing - original draft: E.H., D.D.; Writing - review & editing: E.H., D.D.; Visualization: E.H., C.B., M.-A.B., D.D.; Supervision: E.H., D.D.; Project administration: E.H., D.D.; Funding acquisition: D.D.

## Statistics and reproducibility

Non-parametric tests were used to determine statistical significance, which was set at p values <0.05. Unless stated in figure legend, *in situ* hybridization and immunohistochemistry experiments have been performed on at least 3 limbs of different embryos.

## Data availability

All data are available within the Article and Supplementary Files. Source data for all graphics are provided with this paper. The scRNAseq datasets have been deposited in the NCBI Gene Expression Omnibus database (https://www.ncbi.nlm.nih.gov/geo/) with the accession code numbers GSE166981 (for E4, E6 and E10) and GSE261503 (for E7 and E9). The scRNAseq analysis was performed with standard pipelines, as described in the Methods. Scripts will be made available upon request. The data and detailed protocols in this manuscript are available from the authors upon request.

---

Table S1

Marker list for the 5CT trajectory branches

Excel file of the differentially-expressed genes in each trajectory branch, ordered by decreasing zscore. The ultimate_gene_names have been identified by combined data-mining of the Ensembl and NCBI Entrez databases.

Table S2

Marker list for the nine CT fibroblast clusters at E10

Excel file of the differentially-expressed genes in each cluster, ordered by decreasing fold-change. The ultimate_gene_names have been identified by combined data-mining of the Ensembl and NCBI Entrez databases. Pct.1 is the fraction of marker-expressing cells in the cluster of interest. Pct.2 is the fraction of marker-expressing cells in the other clusters. In the column “cluster” is listed the cluster the marker is associated with for a given row. In the column “cluster list” are listed all the clusters this marker is associated with.

Table S3

Values for the parameters and cut-offs used in the Seurat and the STREAM analyses.

Table S4

List of the primers used for PCR probes

## References

1. Asfour H, Hirsinger E, Rouco R, Zarrouki F, Hayashi S, Swist S, Braun T, Patel K, Relaix F, Andrey G, Stricker S, Duprez D, Stantzou A, Amthor H. (2023) Inhibitory SMAD6 interferes with BMP-dependent generation of muscle progenitor cells and perturbs proximodistal pattern of murine limb muscles. Development. 2023 Jun 1;150(11):dev201504. doi: 10.1242/dev.201504. Epub 2023 Jun 5.

2. Benjamin M, Kaiser E, Milz S. (2008) Structure-function relationships in tendons: a review. J Anat. 2008 Mar;212(3):211-28. doi: 10.1111/j.1469-7580.2008.00864.x.

3. Blitz E, Sharir A, Akiyama H, Zelzer E. (2013) Tendon-bone attachment unit is formed modularly by a distinct pool of Scx- and Sox9-positive progenitors. Development. 2013 Jul;140(13):2680-90. doi: 10.1242/dev.093906. Epub 2013 May 29.

4. Blitz E, Viukov S, Sharir A, Shwartz Y, Galloway JL, Pryce BA, Johnson RL, Tabin CJ, Schweitzer R, Zelzer E. (2009) Bone ridge patterning during musculoskeletal assembly is mediated through SCX regulation of Bmp4 at the tendon-skeleton junction. Dev Cell. 2009 Dec;17(6):861-73. doi: 10.1016/j.devcel.2009.10.010.

5. Bordoni B, Black AC, Varacallo M. (2023) Anatomy, Tendons. 2023 Nov 9. In: StatPearls [Internet]. Treasure Island (FL): StatPearls Publishing; 2024 Jan–. PMID: 30020609

6. Burgy O, Königshoff M. (2018) The WNT signaling pathways in wound healing and fibrosis. Matrix Biol. 2018 Aug;68-69:67-80. doi: 10.1016/j.matbio.2018.03.017. Epub 2018 Mar 20.

7. Chen H, Albergante L, Hsu JY, Lareau CA, Lo Bosco G, Guan J, Zhou S, Gorban AN, Bauer DE, Aryee MJ, Langenau DM, Zinovyev A, Buenrostro JD, Yuan GC, Pinello L. (2019) Single-cell trajectories reconstruction, exploration and mapping of omics data with STREAM. Nat Commun. 2019 Apr 23;10(1):1903. doi: 10.1038/s41467-019-09670-4.

8. Cheng CW, Rifai A, Ka SM, Shui HA, Lin YF, Lee WH, Chen A. (2005) Calcium-binding proteins annexin A2 and S100A6 are sensors of tubular injury and recovery in acute renal failure. Kidney Int. 2005 Dec;68(6):2694-703. doi: 10.1111/j.1523-1755.2005.00740.x.

9. Chevallier, A., Kieny, M. and Mauger, A. (1977). Limb-somite relationship: origin of the limb musculature. J. Embryol. Exp. Morphol. 41, 245–58.

10. Christ, B., Jacob, H. J. and Jacob, M. (1977). Experimental analysis of the origin of the wing musculature in avian embryos. Anat. Embryol. (Berl). 150, 171–86.

11. Colasanto MP, Eyal S, Mohassel P, Bamshad M, Bonnemann CG, Zelzer E, Moon AM, Kardon G (2016) Development of a subset of forelimb muscles and their attachment sites requires the ulnar-mammary syndrome gene Tbx3. Dis Model Mech. 2016 Nov 1;9(11):1257-1269. doi: 10.1242/dmm.025874. Epub 2016 Aug 4.

12. Conway JK, Lex A, Gehlenborg N (2017) UpSetR: an R package for the visualization of intersecting sets and their properties Bioinformatics. 2017 Sep 15;33(18):2938-2940. doi: 10.1093/bioinformatics/btx364.

13. Coreen L., Zaffryar-Eilot S., Kaganovsky A., Odeh A., Hasson P. (2023) Fibroblast diversification is an embryonic process dependent on muscle contraction (back to back paper)

14. De Micheli AJ, Swanson JB, Disser NP, Martinez LM, Walker NR, Oliver DJ, Cosgrove BD, Mendias CL. (2020) Single-cell transcriptomic analysis identifies extensive heterogeneity in the cellular composition of mouse Achilles tendons. Am J Physiol Cell Physiol. 2020 Nov 1;319(5):C885-C894. doi: 10.1152/ajpcell.00372.2020. Epub 2020 Sep 2.

15. Driskell RR, Lichtenberger BM, Hoste E, Kretzschmar K, Simons BD, Charalambous M, Ferron SR, Herault Y, Pavlovic G, Ferguson-Smith AC, Watt FM. (2013). Distinct fibroblast lineages determine dermal architecture in skin development and repair. Nature. 2013 Dec 12;504(7479):277-281. doi: 10.1038/nature12783.

16. Duprez D, Lapointe F, Edom-Vovard F, Kostakopoulou K, Robson L. (1999) Sonic hedgehog (SHH) specifies muscle pattern at tissue and cellular chick level, in the chick limb bud. Mech Dev. 1999 Apr;82(1-2):151-63. doi: 10.1016/s0925-4773(99)00040-4.

17. Duprez, D. (2002). Signals regulating muscle formation in the limb during embryonic development. Int. J. Dev. Biol. 46, 915–926.

18. Dyment NA, Liu CF, Kazemi N, Aschbacher-Smith LE, Kenter K, Breidenbach AP, Shearn JT, Wylie C, Rowe DW, Butler DL. (2013) The paratenon contributes to scleraxis-expressing cells during patellar tendon healing. PLoS One. 2013;8(3):e59944. doi: 10.1371/journal.pone.0059944. Epub 2013 Mar 26.

19. Esteves de Lima, J., Bonnin, M.-A., Birchmeier, C., and Duprez, D. (2016). Muscle contraction is required to maintain the pool of muscle progenitors via YAP and NOTCH during fetal myogenesis. ELife 5, e15593.

20. Esteves de Lima, J., Blavet, C., Bonnin, M.-A., Hirsinger, E., Comai, G., Yvernogeau, L., Delfini, M.-C., Bellenger, L., Mella, S., Nassari, S., et al. (2021). Unexpected contribution of fibroblasts to muscle lineage as a mechanism for limb muscle patterning. Nat. Commun. 12, 3851.

21. Esteves de Lima J, Blavet C, Bonnin MA, Hirsinger E, Havis E, Relaix F, Duprez D. (2022) TMEM8C-mediated fusion is regionalized and regulated by NOTCH signalling during foetal myogenesis. Development. 2022 Jan 15;149(2):dev199928. doi: 10.1242/dev.199928. Epub 2022 Jan 25.

22. Flynn CGK, Ginkel PRV, Hubert KA, Guo Q, Hrycaj SM, McDermott AE, Madruga A, Miller AP, Wellik DM. (2023) Hox11-expressing interstitial cells contribute to adult skeletal muscle at homeostasis. Development. 2023 Feb 15;150(4):dev201026. doi: 10.1242/dev.201026. Epub 2023 Feb 23.

23. Fu W, Yang R, Li J. BMC Biol. (2023) Single-cell and spatial transcriptomics reveal changes in cell heterogeneity during progression of human tendinopathy. 2023 Jun 6;21(1):132. doi: 10.1186/s12915-023-01613-2.

24. Fukada SI, Uezumi A. (2023) Roles and Heterogeneity of Mesenchymal Progenitors in Muscle Homeostasis, Hypertrophy, and Disease. Stem Cells. 2023 Jun 15;41(6):552-559. doi: 10.1093/stmcls/sxad023.

25. Gaut, L. and Duprez, D. (2016). Tendon development and diseases. Wiley Interdiscip. Rev. Dev. Biol. 5, 5–23.

26. Giordani L, He GJ, Negroni E, Sakai H, Law JYC, Siu MM, Wan R, Corneau A, Tajbakhsh S, Cheung TH, Le Grand F. (2019) High-Dimensional Single-Cell Cartography Reveals Novel Skeletal Muscle-Resident Cell Populations. Mol Cell. 2019 May 2;74(3):609-621.e6. doi: 10.1016/j.molcel.2019.02.026. Epub 2019 Mar 25.

27. Giuliani G, Vumbaca S, Fuoco C, Gargioli C, Giorda E, Massacci G, Palma A, Reggio A, Riccio F, Rosina M, Vinci M, Castagnoli L, Cesareni G. (2021). SCA-1 micro-heterogeneity in the fate decision of dystrophic fibro/adipogenic progenitors. Cell Death Dis. 2021 Jan 25;12(1):122. doi: 10.1038/s41419-021-03408-1.

28. Giuliani G, Rosina M, Reggio A. (2022) Signaling pathways regulating the fate of fibro/adipogenic progenitors (FAPs) in skeletal muscle regeneration and disease. FEBS J. 2022 Nov;289(21):6484-6517. doi: 10.1111/febs.16080. Epub 2021 Jul 6.

29. Guerquin MJ, Charvet B, Nourissat G, Havis E, Ronsin O, Bonnin MA, Ruggiu M, Olivera-Martinez I, Robert N, Lu Y, Kadler KE, Baumberger T, Doursounian L, Berenbaum F, Duprez D. (2013) Transcription factor EGR1 directs tendon differentiation and promotes tendon repair. J Clin Invest. 2013 Aug;123(8):3564-76. doi: 10.1172/JCI67521. Epub 2013 Jul 25.

30. Harvey T, Flamenco S, Fan CM. (2019) A Tppp3^+^Pdgfra^+^ tendon stem cell population contributes to regeneration and reveals a shared role for PDGF signalling in regeneration and fibrosis. Nat Cell Biol. 2019 Dec;21(12):1490-1503. doi: 10.1038/s41556-019-0417-z. Epub 2019 Nov 25.

31. Hasson, P., DeLaurier, A., Bennett, M., Grigorieva, E., Naiche, L. A., Papaioannou, V. E., Mohun, T. J. and Logan, M. P. O. (2010). Tbx4 and Tbx5 acting in connective tissue are required for limb muscle and tendon patterning. Dev. Cell 18, 148–56.

32. Havis, E., Bonnin, M. A., Olivera-Martinez, I., Nazaret, N., Ruggiu, M., Weibel, J., Durand, C., Guerquin, M.-J., Bonod-Bidaud, C., Ruggiero, F., et al. (2014). Transcriptomic analysis of mouse limb tendon cells during development. Development 141, 3683–96.

33. Havis E, Bonnin MA, Esteves de Lima J, Charvet B, Milet C, Duprez D. (2016) TGFβ and FGF promote tendon progenitor fate and act downstream of muscle contraction to regulate tendon differentiation during chick limb development. Development. 2016 Oct 15;143(20):3839-3851. doi: 10.1242/dev.136242. Epub 2016 Sep 13.

34. Helle KB. (2010) Chromogranins A and B and secretogranin II as prohormones for regulatory peptides from the diffuse neuroendocrine system. Results Probl Cell Differ. 2010;50:21–44. doi: 10.1007/400_2009_26.

35. Helmbacher F, Stricker S. (2020) Tissue cross talks governing limb muscle development and regeneration. Semin Cell Dev Biol. 2020 Aug;104:14-30. doi: 10.1016/j.semcdb.2020.05.005. Epub 2020 Jun 7.

36. Hogarth MW, Defour A, Lazarski C, Gallardo E, Diaz Manera J, Partridge TA, Nagaraju K, Jaiswal JK. (2019) Fibroadipogenic progenitors are responsible for muscle loss in limb girdle muscular dystrophy 2B. Nat Commun. 2019 Jun 3;10(1):2430. doi: 10.1038/s41467-019-10438-z.

37. Hornik C, Krishan K, Yusuf F, Scaal M, Brand-Saberi B. (2005) cDermo-1 misexpression induces dense dermis, feathers, and scales. Dev Biol. 2005 Jan 1;277(1):42-50. doi: 10.1016/j.ydbio.2004.08.050.

38. Huang DW, Sherman BT, Lempicki RA. (2009) Systematic and integrative analysis of large gene lists using DAVID bioinformatics resources. Nat Protoc. 2009;4(1):44-57. doi: 10.1038/nprot.2008.211.

39. Izu Y, Adams SM, Connizzo BK, Beason DP, Soslowsky LJ, Koch M, Birk DE. (2021) Collagen XII mediated cellular and extracellular mechanisms regulate establishment of tendon structure and function. Matrix Biol. 2021 Jan;95:52-67. doi: 10.1016/j.matbio.2020.10.004. Epub 2020 Oct 20.

40. Kardon, G. (1998). Muscle and tendon morphogenesis in the avian hind limb. Development 125, 4019–32.

41. Kardon, G., Harfe, B. D. and Tabin, C. J. (2003). A Tcf4-positive mesodermal population provides a prepattern for vertebrate limb muscle patterning. Dev. Cell 5, 937–44.

42. Kardon G. (2011) Development of the musculoskeletal system: meeting the neighbors. Development. 2011 Jul;138(14):2855-9. doi: 10.1242/dev.067181.

43. Kawano Y, Yoshimura T, Kaibuchi K. Nihon Yakurigaku Zasshi. (2002) [Involvement of small GTPase Rho in cardiovascular diseases]. 2002 Sep;120(3):149-58. doi: 10.1254/fpj.120.149.

44. Kendal AR, Layton T, Al-Mossawi H, Appleton L, Dakin S, Brown R, Loizou C, Rogers M, Sharp R, Carr A. (2020) Multi-omic single cell analysis resolves novel stromal cell populations in healthy and diseased human tendon. Sci Rep. 2020 Sep 3;10(1):13939. doi: 10.1038/s41598-020-70786-5.

45. Kotsaris G, Qazi TH, Bucher CH, Zahid H, Pöhle-Kronawitter S, Ugorets V, Jarassier W, Börno S, Timmermann B, Giesecke-Thiel C, Economides AN, Le Grand F, Vallecillo-García P, Knaus P, Geissler S, Stricker S. (2023) Odd skipped-related 1 controls the pro-regenerative response of fibro-adipogenic progenitors. NPJ Regen Med. 2023 Apr 5;8(1):19. doi: 10.1038/s41536-023-00291-6.

46. Kristensen LS, Raynor MP, Candiloro I, Dobrovic A. Oncotarget. (2012) Methylation profiling of normal individuals reveals mosaic promoter methylation of cancer-associated genes. 2012 Apr;3(4):450-61. doi: 10.18632/oncotarget.480.

47. EGR1 and EGR2 involvement in vertebrate tendon differentiation. (2011) Lejard V, Blais F, Guerquin MJ, Bonnet A, Bonnin MA, Havis E, Malbouyres M, Bidaud CB, Maro G, Gilardi-Hebenstreit P, Rossert J, Ruggiero F, Duprez D. J Biol Chem. 2011 Feb 18;286(7):5855-67. doi: 10.1074/jbc.M110.153106. Epub 2010 Dec 20.

48. Li L, Cserjesi P, Olson EN. (1995) Dermo-1: a novel twist-related bHLH protein expressed in the developing dermis. Dev Biol. 1995 Nov;172(1):280-92. doi:10.1006/dbio.1995.0023.

49. Li T, Xiong X, Wang Y, Li Y, Liu Y, Zhang M, Li C, Yu T, Cao W, Chen S, Zhang H, Wang X, Lv L, Zhou Y, Liang H, Li X, Shan H. (2023) Neuroepithelial cell-transforming 1 promotes cardiac fibrosis via the Wnt/β-catenin signaling pathway. iScience. 2023 Sep 9;26(10):107888. doi: 10.1016/j.isci.2023.107888. eCollection 2023 Oct 20.

50. Lim HI, Hajjar KA. (2021) Annexin A2 in Fibrinolysis, Inflammation and Fibrosis. Int J Mol Sci. 2021 Jun 25;22(13):6836. doi: 10.3390/ijms22136836.

51. Lynch MD, Watt FM. (2018) Fibroblast heterogeneity: implications for human disease. J Clin Invest. 2018 Jan 2;128(1):26-35. doi: 10.1172/JCI93555. Epub 2018 Jan 2.

52. Macosko EZ, Basu A, Satija R, Nemesh J, Shekhar K, Goldman M, Tirosh I, Bialas AR, Kamitaki N, Martersteck EM, Trombetta JJ, Weitz DA, Sanes JR, Shalek AK, Regev A, McCarroll SA. (2015) Highly Parallel Genome-wide Expression Profiling of Individual Cells Using Nanoliter Droplets. Cell. 2015 May 21;161(5):1202-1214. doi: 10.1016/j.cell.2015.05.002.

53. Martinerie C, Garcia M, Do TT, Antoine B, Moldes M, Dorothee G, Kazazian C, Auclair M, Buyse M, Ledent T, Marchal PO, Fesatidou M, Beisseiche A, Koseki H, Hiraoka S, Chadjichristos CE, Blondeau B, Denis RG, Luquet S, Fève B. (2016) NOV/CCN3: A New Adipocytokine Involved in Obesity-Associated Insulin Resistance. Diabetes. 2016 Sep;65(9):2502-15. doi: 10.2337/db15-0617. Epub 2016 Jun 9.

54. Mathew, S. J., Hansen, J. M., Merrell, A. J., Murphy, M. M., Lawson, J. a, Hutcheson, D. a, Hansen, M. S., Angus-Hill, M. and Kardon, G. (2011). Connective tissue fibroblasts and Tcf4 regulate myogenesis. Development 138, 371–84.

55. McQueen C, Towers M. Development. (2020) Establishing the pattern of the vertebrate limb. 2020 Sep 11;147(17):dev177956. doi: 10.1242/dev.177956.

56. Méndez-Barbero N, San Sebastian-Jaraba I, Blázquez-Serra R, Martín-Ventura JL, Blanco-Colio LM. (2022) Annexins and cardiovascular diseases: Beyond membrane trafficking and repair. Front Cell Dev Biol. 2022 Oct 14;10:1000760. doi: 10.3389/fcell.2022.1000760. eCollection 2022.

57. Mercader N, Leonardo E, Piedra ME, Martínez-A C, Ros MA, Torres M. (2000) Opposing RA and FGF signals control proximodistal vertebrate limb development through regulation of Meis genes. Development. 2000 Sep;127(18):3961-70. doi: 10.1242/dev.127.18.3961.

58. Mienaltowski MJ, Adams SM, Birk DE. (2013) Regional differences in stem cell/progenitor cell populations from the mouse achilles tendon. Tissue Eng Part A. 2013 Jan;19(1-2):199-210. doi: 10.1089/ten.TEA.2012.0182. Epub 2012 Sep 14.

59. Mienaltowski, M. J., Adams, S. M. and Birk, D. E. (2014). Tendon proper- and peritenon- derived progenitor cells have unique tenogenic properties. Stem Cell Res. Ther. 5, 86.

60. Mienaltowski MJ, Cánovas A, Fates VA, Hampton AR, Pechanec MY, Islas-Trejo A, Medrano JF. (2019) Transcriptome profiles of isolated murine Achilles tendon proper- and peritenon-derived progenitor cells. J Orthop Res. 2019 Jun;37(6):1409-1418. doi: 10.1002/jor.24076. Epub 2018 Jul 13.

61. Murchison, N. D., Price, B. A., Conner, D. A., Keene, D. R., Olson, E. N., Tabin, C. J. and Schweitzer, R. (2007). Regulation of tendon differentiation by scleraxis distinguishes force-transmitting tendons from muscle-anchoring tendons. Development 134, 2697–708.

62. Murphy MM, Lawson JA, Mathew SJ, Hutcheson DA, Kardon G. (2011) Satellite cells, connective tissue fibroblasts and their interactions are crucial for muscle regeneration. Development. 2011 Sep;138(17):3625-37. doi: 10.1242/dev.064162.

63. Nassari, S., Duprez, D. and Fournier-Thibault, C. (2017). Non-myogenic contribution to muscle development and homeostasis: the role of connective tissues. Front. Cell Dev. Biol. 5, 22.

64. Nelson CE, Morgan BA, Burke AC, Laufer E, DiMambro E, Murtaugh LC, Gonzales E, Tessarollo L, Parada LF, Tabin C. (1996) Analysis of Hox gene expression in the chick limb bud. Development. 1996 May;122(5):1449-66. doi: 10.1242/dev.122.5.1449.

65. Nourissat G, Berenbaum F, Duprez D. (2015) Tendon injury: from biology to tendon repair. Nat Rev Rheumatol. 2015 Apr;11(4):223-33. doi: 10.1038/nrrheum.2015.26. Epub 2015 Mar 3.

66. Orgeur M, Martens M, Leonte G, Nassari S, Bonnin MA, Börno ST, Timmermann B, Hecht J, Duprez D, Stricker S. (2018) Genome-wide strategies identify downstream target genes of chick connective tissue-associated transcription factors. Development. 2018 Mar 29;145(7):dev161208. doi: 10.1242/dev.161208.

67. Ono Y, Schlesinger S, Fukunaga K, Yambe S, Sato T, Sasaki T, Shukunami C, Asahara H, Inui M. (2023) Scleraxis-lineage cells are required for correct muscle patterning. Development. 2023 May 15;150(10):dev201101. doi: 10.1242/dev.201101. Epub 2023 May 29.

68. Pascal A, Gallaud E, Giet R, Benaud C. (2022) Annexin A2 and Ahnak control cortical NuMA-dynein localization and mitotic spindle orientation. J Cell Sci. 2022 May 1;135(9):jcs259344. doi: 10.1242/jcs.259344. Epub 2022 May 6.

69. Pechanec MY, Mienaltowski MJ. (2023) Decoding the transcriptomic expression and genomic methylation patterns in the tendon proper and its peritenon region in the aging horse. BMC Res Notes. 2023 Oct 11;16(1):267. doi: 10.1186/s13104-023-06562-1.

70. Pena-Amaro (2021). The musculotendinous transition of the extracellular matrix. Apunts Sport medicin. April-June 2021 Vol. 56. Issue 210 DOI: 10.1016/j.apunsm.2021.100350 Vol. 56. Issue 210

71. Pineault KM, Wellik DM. (2014) Hox genes and limb musculoskeletal development. Curr Osteoporos Rep. 2014 Dec;12(4):420-7. doi: 10.1007/s11914-014-0241-0.

72. Purslow PP (2020) The Structure and Role of Intramuscular Connective Tissue in Muscle Function. Front Physiol. 2020 May 19;11:495. doi: 10.3389/fphys.2020.00495. eCollection 2020.

73. Raajendiran A, Ooi G, Bayliss J, O’Brien PE, Schittenhelm RB, Clark AK, Taylor RA, Rodeheffer MS, Burton PR, Watt MJ. (2019) Identification of Metabolically Distinct Adipocyte Progenitor Cells in Human Adipose Tissues. Cell Rep. 2019 Apr 30;27(5):1528- 1540.e7. doi: 10.1016/j.celrep.2019.04.010.

74. Rolnick KI, Choe JA, Leiferman EM, Kondratko-Mittnacht J, Clements AEB, Baer GS, Jiang P, Vanderby R, Chamberlain CS. (2022) Periostin modulates extracellular matrix behavior in tendons. Matrix Biol Plus. 2022 Nov 9;16:100124. doi: 10.1016/j.mbplus.2022.100124. eCollection 2022 Dec.

75. Scott RW, Arostegui M, Schweitzer R, Rossi FMV, Underhill TM. (2019) Hic1 Defines Quiescent Mesenchymal Progenitor Subpopulations with Distinct Functions and Fates in Skeletal Muscle Regeneration. Cell Stem Cell. 2019 Dec 5;25(6):797-813.e9. doi: 10.1016/j.stem.2019.11.004.

76. Sefton EM, Kardon G. (2019) Connecting muscle development, birth defects, and evolution: An essential role for muscle connective tissue. Curr Top Dev Biol. 2019;132:137–176. doi: 10.1016/bs.ctdb.2018.12.004. Epub 2019 Jan 3.

77. Sherman BT, Hao M, Qiu J, Jiao X, Baseler MW, Lane HC, Imamichi T, Chang W. (2022) DAVID: a web server for functional enrichment analysis and functional annotation of gene lists (2021 update). Nucleic Acids Res. 2022 Jul 5;50(W1):W216-W221. doi: 10.1093/nar/gkac194.

78. Shukunami C, Takimoto A, Oro M, Hiraki Y. (2006) Scleraxis positively regulates the expression of tenomodulin, a differentiation marker of tenocytes. Dev Biol. 2006 Oct 1;298(1):234-47. doi: 10.1016/j.ydbio.2006.06.036. Epub 2006 Jun 27.

79. Shukunami C, Takimoto A, Nishizaki Y, Yoshimoto Y, Tanaka S, Miura S, Watanabe H, Sakuma T, Yamamoto T, Kondoh G, Hiraki Y. (2018) Scleraxis is a transcriptional activator that regulates the expression of Tenomodulin, a marker of mature tenocytes and ligamentocytes. Sci Rep. 2018 Feb 16;8(1):3155. doi: 10.1038/s41598-018-21194-3.

80. Steffen D, Mienaltowski M, Baar K. (2023) Spatial gene expression in the adult rat patellar tendon. Matrix Biol Plus. 2023 Nov 25;19-20:100138. doi: 10.1016/j.mbplus.2023.100138. eCollection 2023 Dec.

81. Stricker, S., Mathia, S., Haupt, J., Seemann, P., Meier, J. and Mundlos, S. (2012). Odd-skipped related genes regulate differentiation of embryonic limb mesenchyme and bone marrow mesenchymal stromal cells. Stem Cells Dev. 21, 623–33.

82. Stumm J, Vallecillo-García P, Vom Hofe-Schneider S, Ollitrault D, Schrewe H, Economides AN, Marazzi G, Sassoon DA, Stricker S. (2018) Odd skipped-related 1 (Osr1) identifies muscle-interstitial fibro-adipogenic progenitors (FAPs) activated by acute injury. Stem Cell Res. 2018 Oct;32:8-16. doi: 10.1016/j.scr.2018.08.010. Epub 2018 Aug 10.

83. Swinehart IT, Schlientz AJ, Quintanilla CA, Mortlock DP, Wellik DM. (2013) Hox11 genes are required for regional patterning and integration of muscle, tendon and bone. Development. 2013 Nov;140(22):4574-82. doi: 10.1242/dev.096693. Epub 2013 Oct 23.

84. ten Berge D, Brouwer A, Korving J, Martin JF, Meijlink F. (1998) Prx1 and Prx2 in skeletogenesis: roles in the craniofacial region, inner ear and limbs. Development. 1998 Oct;125(19):3831-42. doi: 10.1242/dev.125.19.3831.

85. Vallecillo García, P., Orgeur, M., Vom Hofe-Schneider, S., Stumm, J., Kappert, V., Ibrahim, D. M., Börno, S. T., Hayashi, S., Relaix, F., Hildebrandt, K., et al. (2017). Odd skipped-related 1 identifies a population of embryonic fibro-adipogenic progenitors regulating myogenesis during limb development. Nat. Commun. 8, 1218

86. Vorstandlechner V, Laggner M, Kalinina P, Haslik W, Radtke C, Shaw L, Lichtenberger BM, Tschachler E, Ankersmit HJ, Mildner M. (2020) Deciphering the functional heterogeneity of skin fibroblasts using single-cell RNA sequencing. FASEB J. 2020 Mar;34(3):3677-3692. doi: 10.1096/fj.201902001RR. Epub 2020 Jan 12.

87. Wang S, Yu J, Kane MA, Moise AR. (2020) Modulation of retinoid signaling: therapeutic opportunities in organ fibrosis and repair. Pharmacol Ther. 2020 Jan;205:107415. doi: 10.1016/j.pharmthera.2019.107415. Epub 2019 Oct 16.

88. Wang T, Wagner A, Gehwolf R, Yan W, Passini FS, Thien C, Weissenbacher N, Lin Z, Lehner C, Teng H, Wittner C, Zheng Q, Dai J, Ni M, Wang A, Papadimitriou J, Leys T, Tuan RS, Senck S, Snedeker JG, Tempfer H, Jiang Q, Zheng MH, Traweger A. (2021) Load-induced regulation of tendon homeostasis by SPARC, a genetic predisposition factor for tendon and ligament injuries. Sci Transl Med. 2021 Feb 24;13(582):eabe5738. doi: 10.1126/scitranslmed.abe5738.

89. Wickham, H. (2016). ggplot2: Elegant Graphics for Data Analysis. New York, NY: Springer.

90. Wilmerding, A., Rinaldi, L., Caruso, N., Lo Re, L., Bonzom, E., Saurin, A.J., Graba, Y., and Delfini, M.-C. (2021). HoxB genes regulate neuronal delamination in the trunk neural tube by controlling the expression of Lzts1. Development 148, dev195404.

91. Wolock, S. L., Lopez, R. & Klein, A. M. Scrublet: computational identification of cell doublets in single-cell transcriptomic data. Cell Syst. 8, 281–291.e9. 2019.

92. Xing E, Ma F, Wasikowski R, Billi AC, Gharaee-Kermani M, Fox J, Dobry C, Victory A, Sarkar MK, Xing X, Plazyo O, Chen HW, Barber G, Jacobe H, Tsou PS, Modlin RL, Varga J, Kahlenberg JM, Tsoi LC, Gudjonsson JE, Khanna D. (2023) Pansclerotic morphea is characterized by IFN-γ responses priming dendritic cell fibroblast crosstalk to promote fibrosis. JCI Insight. 2023 Aug 22;8(16):e171307. doi: 10.1172/jci.insight.171307.

93. Yan R, Zhang H, Ma Y, Lin R, Zhou B, Zhang T, Fan C, Zhang Y, Wang Z, Fang T, Yin Z, Cai Y, Ouyang H, Chen X. (2022) Discovery of Muscle-Tendon Progenitor Subpopulation in Human Myotendinous Junction at Single-Cell Resolution. Research (Wash D C). 2022 Sep 28;2022:9760390. doi: 10.34133/2022/9760390. eCollection 2022.

94. Yaseen, W., Kraft-Sheleg, O., Zaffryar-Eilot, S., Melamed, S., Sun, C., Millay, D.P., and Hasson, P. (2021). Fibroblast fusion to the muscle fiber regulates myotendinous junction formation. Nat. Commun. 12, 3852.

95. Yin Z, Hu JJ, Yang L, Zheng ZF, An CR, Wu BB, Zhang C, Shen WL, Liu HH, Chen JL, Heng BC, Guo GJ, Chen X, Ouyang HW. (2016) Single-cell analysis reveals a nestin^+^ tendon stem/progenitor cell population with strong tenogenic potentiality. Sci Adv. 2016 Nov 18;2(11):e1600874. doi: 10.1126/sciadv.1600874. eCollection 2016 Nov.

96. Zappia, L. & Oshlack, A. Clustering trees: a visualization for evaluating clusterings at multiple resolutions. Gigascience 7, giy083 (2018).

97. Zhao L, Son JS, Wang B, Tian Q, Chen Y, Liu X, de Avila JM, Zhu MJ, Du M. (2020) Retinoic acid signalling in fibro/adipogenic progenitors robustly enhances muscle regeneration. EBioMedicine. 2020 Oct;60:103020. doi: 10.1016/j.ebiom.2020.103020. Epub 2020 Sep 24.

98. Zhu M, Metzen F, Hopkinson M, Betz J, Heilig J, Sodhi J, Imhof T, Niehoff A, Birk DE, Izu Y, Krüger M, Pitsillides AA, Altmüller J, van Osch GJVM, Straub V, Schreiber G, Paulsson M, Koch M, Brachvogel B. (2023) Ablation of collagen XII disturbs joint extracellular matrix organization and causes patellar subluxation. iScience. 2023 Jun 28;26(7):107225. doi: 10.1016/j.isci.2023.107225. eCollection 2023 Jul 21.

