## Supplemental Figures for "Limb connective tissue is organized in a continuum of promiscuous fibroblast identities during development"

Estelle Hirsinger et al.

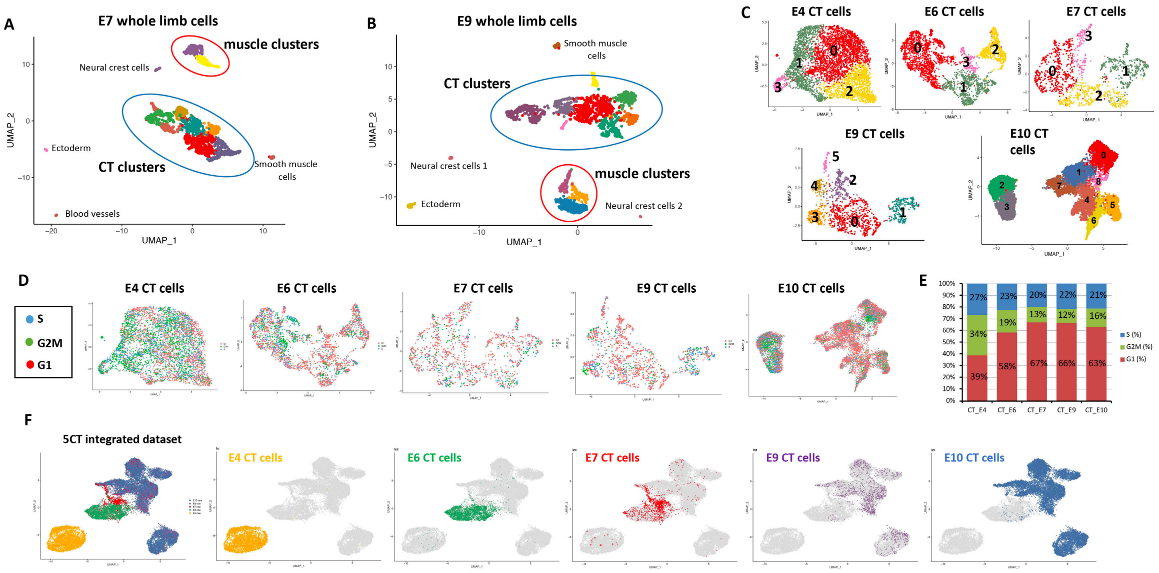


**Figure S1**

**Clustering of whole limb and CT datasets**

(**A,B**) UMAP plots showing the cluster distribution of whole limb E7 (1546 cells) (A) and E9 (1893 cells) (B) datasets. CT and muscle clusters are circled. **(C)** UMAP plots showing the distribution of CT clusters in E4 (4654 cells), E6 (2869 cells), E7 (1229 cells), E9 (1328 cells) and E10 (14490 cells) CT datasets. (**D**) UMAP plots showing the distribution of the cell cycle phases (S, G2/M and G1) in E4, E6, E7, E9 and E10 CT datasets. (**E**) Barplot showing the *in silico* quantification of the number of cells in each cell cycle phase (S, G2/M and G1) at successive developmental stages. (**F**) UMAP plots showing the distribution of the datasets of origin in the 5CT integrated dataset. The E4 (yellow), E6 (green), E7 (red), E9 (purple) and E10 (bleu) original CT datasets were plotted together (left panel, same as Figure 1A, shown again for clarity) or individually.

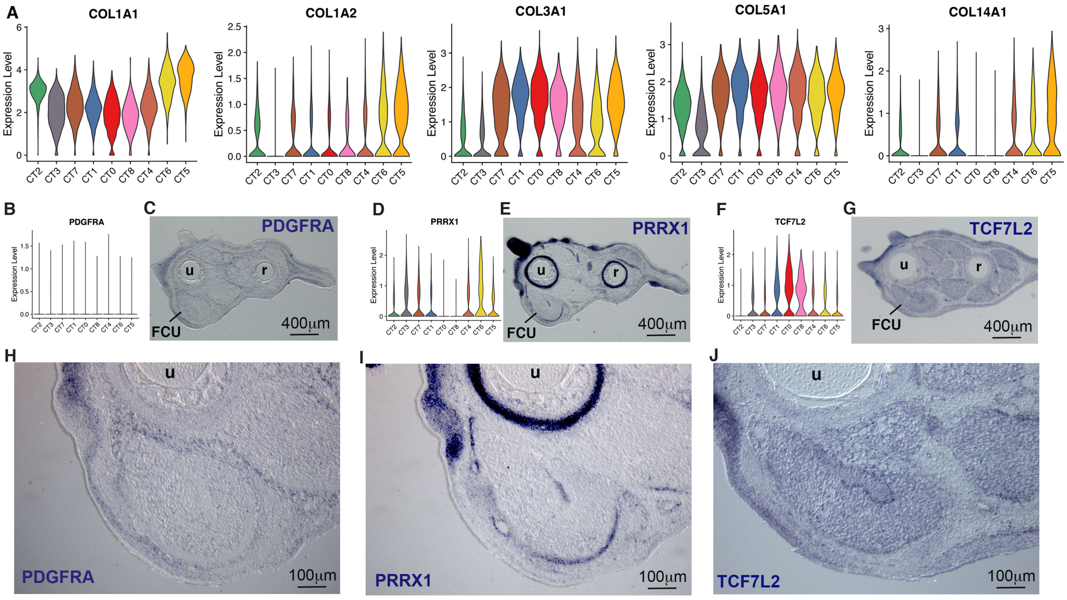


**Figure S2**

**Expression of generic CT markers in fibroblast clusters and limbs**

(**A**) Violin plots showing the log-normalized expression levels across clusters for the *COL1A1, COL1A2, COL3A1*, *COL5A1* and *COL14A1* genes. (**B,D,F**) Violin plots showing the log-normalized expression levels across clusters for the *PDFGRA* (B), *PRRX1* (D) and *TCF7L2* (F) genes. (**C,E,G,H,I,J**) Colorimetric *in situ* hybridization to transverse limb sections of E10 chicken embryos with PDGFRA (C,H), PRRX1 (E,I) and TCF7L2 (G,J) probes (blue). (H,I,J) show high magnifications of the FCU muscle from limb sections (C,E,G).

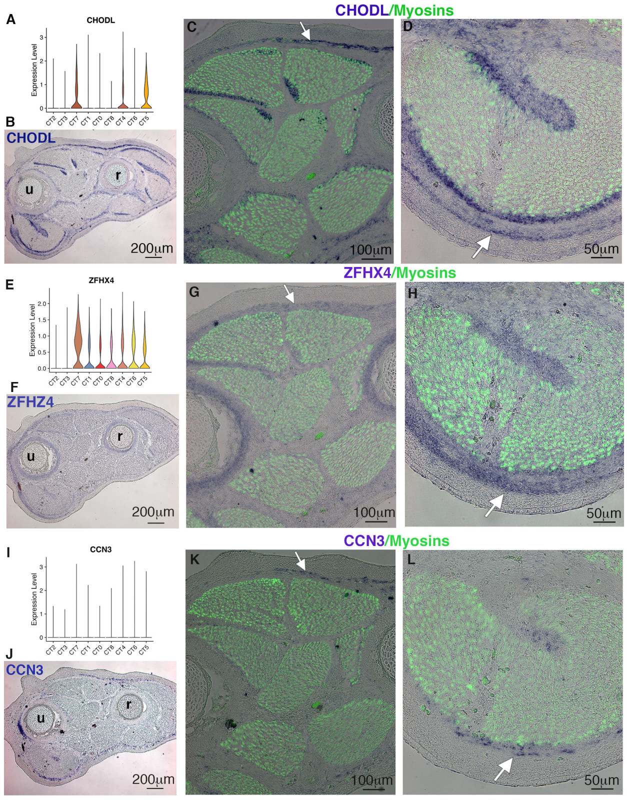


M

| | Cell number in cluster 7 | % vs cluster 7 |
|---|---|---|
| Cluster 7 | 1088 | 100% |
| ZFHX4+ cells | 746 | 69% |
| CHODL+ cells | 450 | 41% |
| CCN3+ cells | 221 | 20% |
| Any combinations ZFHX4/CHODL/CCN3 | 915 | 84% |

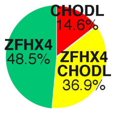


**Figure S3**

**Expression of markers for MCT cluster 7 fibroblasts**

(**A**) Violin plot showing the log-normalized expression levels across clusters for the *CHODL* gene. Panel also shown in Figure 5A. (**B**) Colorimetric *in situ* hybridization to transverse limb sections of E10 chicken embryos with CHODL probe (blue labeling). (**C,D**) High magnification of dorsal limb regions (C) and FCU muscle (D) combined with myosin location with immunostaining with MF20 antibody (green). White arrows show the CHODL expression in the cluster 7 fibroblast layer. (**E**) Violin plot showing the log-normalized expression levels across clusters for the *ZFHX4* gene. (**F**) *In situ* hybridization to transverse limb sections of E10 chicken embryos with ZFHX4 probe (blue labeling). (**G,H**) High magnification of dorsal limb regions (G) and FCU muscle (H) combined with myosin location with immunostaining with MF20 antibody (green). White arrows show the *ZFHX4* expression in cluster 7 fibroblast layer. (**I**) Violin plot showing the log-normalized expression levels across clusters for the *CCN3* gene. (**J**) *In situ* hybridization to transverse limb sections of E10 chicken embryos with CCN3 probe (blue labeling). (**K,L**) High magnification of dorsal limb regions (K) and FCU muscle (L) combined with myosin location, immunostaining with MF20 antibody (green). White arrows show the *CCN3* expression in cluster 7 fibroblast layer. (C,D,G,H,K,L) White arrows show the cluster 7 fibroblast layer. (**M**) Number of *ZFHX4+, CHODL+, CCN3+* fibroblasts in cluster 7. (**N**) Percentage of *CHODL+, ZFHX4+* and *CHODL+/ZFHX4+* fibroblasts among the *CHODL-ZFHX4* population within cluster 7.

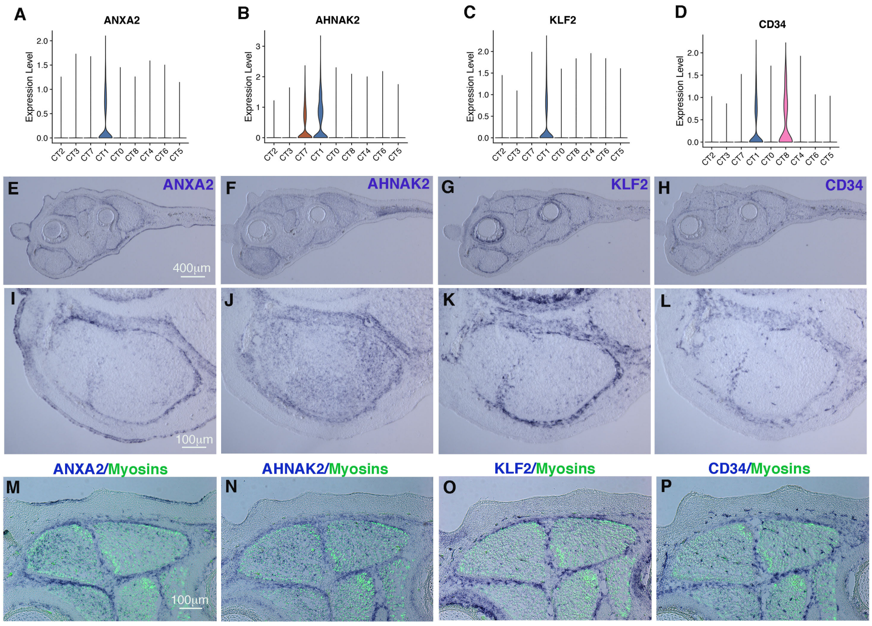


Q

| | Cell number in cluster 1 | % vs cluster 1 |
|---|---|---|
| Cluster 1 | 2294 | 100% |
| ANXA2+ cells | 748 | 33% |
| AHNAK2+ cells | 1198 | 52% |
| KLF2+ cells | 719 | 31% |
| CD34+ cells | 800 | 35% |
| any combinations ANXA2/AHNAK2/KLF2/CD34 | 1834 | 80% |

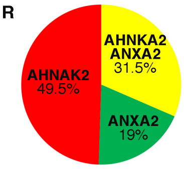


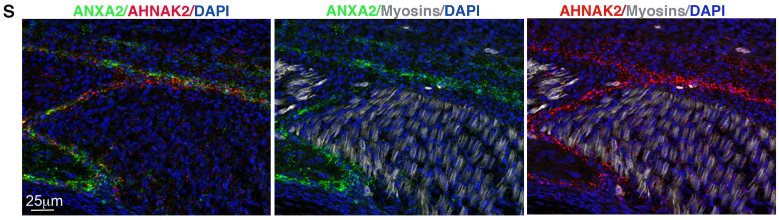


**Figure S4**

**Expression of markers for MCT cluster 1 fibroblasts**

(**A-D**) Violin plots showing the log-normalized expression levels across clusters for representative markers of cluster 1, the *ANXA2* (A, panel also shown in Figure 5B), *AHNAK2* (B), *KLF2* (C), *CD34* (D) genes. (**E-L**) Colorimetric *in situ* hybridization with ANXA2 (E,I), AHNAK2 (F,J), KLF2 (G,K), CD34 (H,L) probes (blue labeling) to transverse limb sections from E10 chicken embryos. (I-L) are high magnification of FCU muscles of limbs shown in (E-H). (**M-P**) *In situ* hybridization to transverse limb sections with ANXA2 (M), AHNAK2 (N), KLF2 (O), CD34 (P) probes (blue labeling) focused on dorsal limb regions and combined with immunostaining with the MF20 antibody to visualize muscle (green). (**Q**) Numbers and percentage of *ANXA2+, AHNAK2+, KLF2+, CD34+* fibroblasts in cluster 1. (**R**) Percentage of *ANXA2+, AHNAK2+* and double *ANXA2+/AHNAK2+* fibroblasts among the *ANXA2-AHNAK2* population within cluster 1. (**S**) Double fluorescent *in situ* hybridization to transverse limb sections of E10 chicken embryos, focused on a dorsal muscle, with ANXA2 (green) and AHNAK2 (red) probes followed with immunostaining with the MF20 antibody to label myosins (grey).

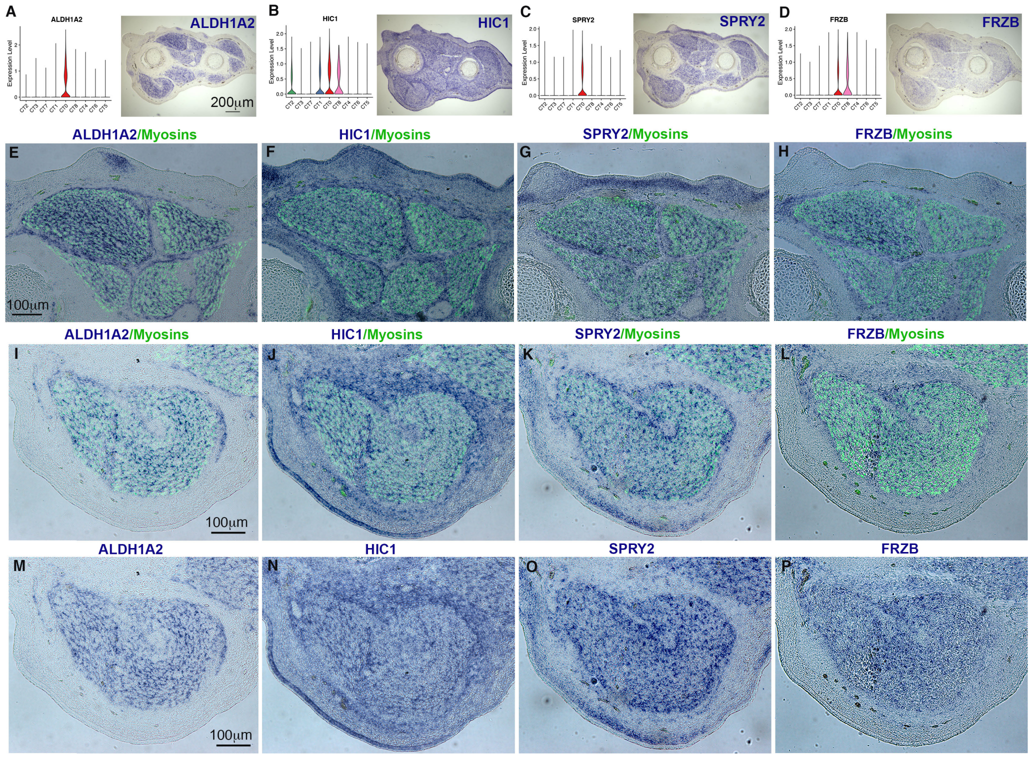


Q

| | Cell number in cluster 0 | Percentage vs cluster 0 |
|---|---|---|
| Cluster 0 | 2745 | 100% |
| ALDH1A2+ cells | 1417 | 52% |
| HIC1+ cells | 1408 | 51% |
| SPRY2+ cells | 1006 | 37% |
| FRZB1+ cells | 907 | 33% |
| ALDH1A2/HIC1/SPRY2/FRZB1 combinations | 2418 | 88% |

R

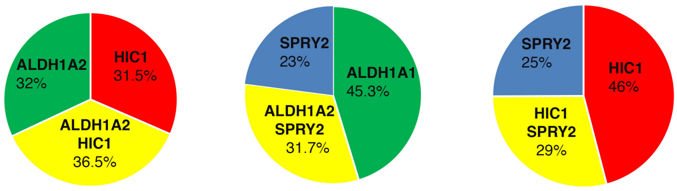


### Figure S5

### Expression of markers for MCT cluster 0 fibroblasts

(**A-D**) Expression of representative genes of cluster 0, *ALDH1A2* (A), *HIC1* (B), *SPRY2* (C) and *FRZB* (D) with violin plots and *in situ* hybridization to transverse limb sections with corresponding probes. The violin plot for *ALDH1A2* is also shown in Figure 5C. (**E-H**) Colorimetric *in situ* hybridization to transverse limb sections with ALDH1A2 (E), HIC1 (F), SPRY2 (G) and FRZB (H) probes (blue labeling) followed by an immunohistochemistry with the MF20 antibody to label myosins (green), focused on dorsal limb regions. (**I-P**) Colorimetric *in situ* hybridization to transverse limb sections with ALDH1A2 (I), HIC1 (J), SPRY2 (K) and FRZB (L) probes (blue labeling) followed by an immunohistochemistry with the MF20 antibody to label myosins (green), focused on the FCU muscle. (M-P) are the same photos of (I-L) without the MF20 staining. (**Q**) Cell numbers and percentage for *ALDH1A2+, HIC1+, SPRY2+, FRZB1+* fibroblasts in cluster 0. (**R**) Percentage of *ALDH1A2+, HIC1+* and double *ALDH1A2+/HIC1+* fibroblasts among the *ALDH1A2-HIC1* population within cluster 0. Percentage of *ALDH1A2+, SPRY2+* and double *ALDH1A2+/SPRY2+* fibroblasts among the *ALDH1A2-SPRY2* population within cluster 0. Percentage of *HIC1+, SPRY2+* and double *HIC1+/ SPRY2+* fibroblasts among the *HIC1-SPRY2* population within cluster 0.

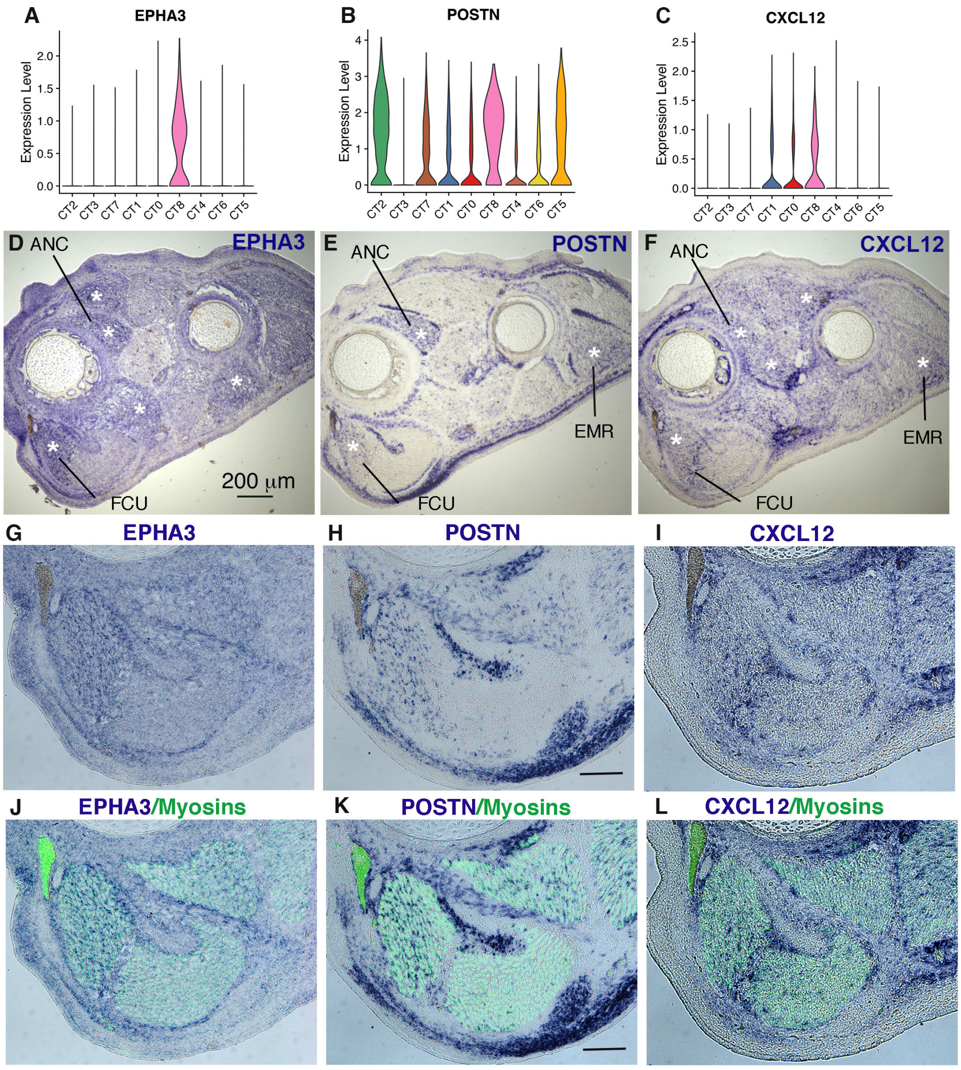


M

| | Cell number in cluster 8 | % vs cluster 8 |
|---|---|---|
| Cluster 8 | 308 | 100% |
| POSTN+ cells | 245 | 80% |
| EPHA3+ cells | 194 | 63% |
| CXCL12+ cells | 135 | 44% |
| Any combinations POSTN/EPHA3/CXCL12 | 289 | 94% |

N

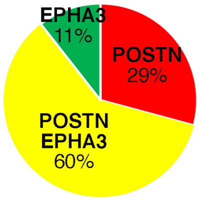


**Figure S6**
**Expression of markers for MCT cluster 8 fibroblasts**
(**A-C**) Violin plots showing the log-normalized expression levels across clusters for representative markers of cluster 8, *EPHA3* (A, shown in Figure 5D), *POSTN* (B) and *CXCL12* (C). (**D-L**) Colorimetric *in situ* hybridization to transverse limb sections of E10 chicken embryos. (G,H,I) show the FCU muscles of limb sections (D,E,F) hybridized with EPHA3 (D,G), POSTN (E,H) and CXCL12 (F,I) probes. (J,K,L) show the same FCU muscle combined with immnunostaining with the MF20 antibody of (G,H,I). (**M**) Cell numbers and percentage of *EPHA3+*, *POSTN+*, *CXCL12+* fibroblasts in cluster 8. (**N**) Percentage of *EPHA3+*, *POSTN+* and double *EPHA3+/POSTN+* fibroblasts among the *EPHA3-POSTN* population within cluster 8.

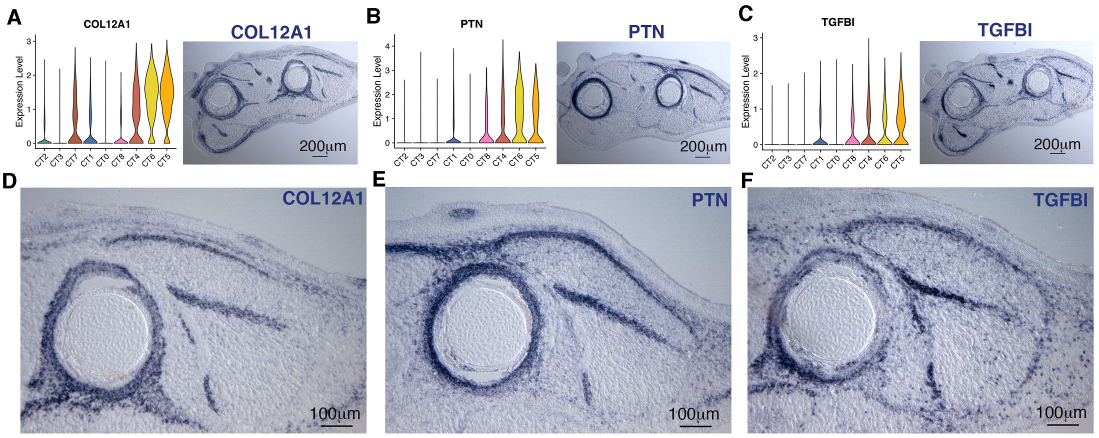


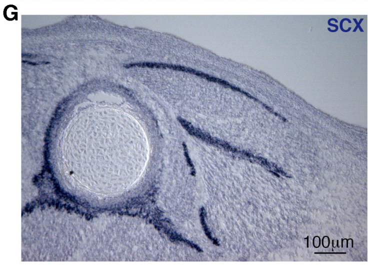


H

| | Cell number in clusters 4,5,6 | % vs clusters 4,5,6 |
|---|---|---|
| Clusters 4,5,6 | 4191 | 100% |
| COL12A1+ cells | 3382 | 81% |
| PTN+ cells | 2391 | 57% |
| TGFBI+ cells | 2163 | 52% |
| COL12A1+/PTN+/TGFBI+ cells | 1162 | 30% |
| Any combinations COL12A1/PTN/TGFBI | 3911 | 93% |

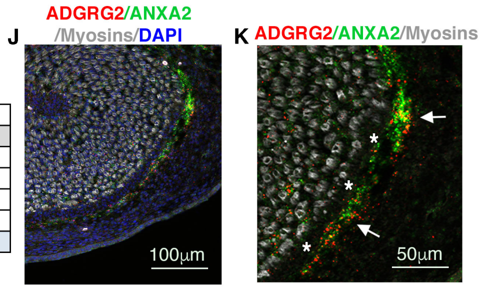


I

| | Cell number in cluster 4 | % vs cluster 4 |
|---|---|---|
| Cluster 4 | 1729 | 100% |
| COL12A1+ cells | 1163 | 67% |
| PTN+ cells | 799 | 46% |
| TGFBI+ cells | 768 | 44% |
| ADGRG2+ cells | 582 | 34% |
| Any combinations COL12A1/PTN/TGFBI/ADGRG2 | 1573 | 91% |

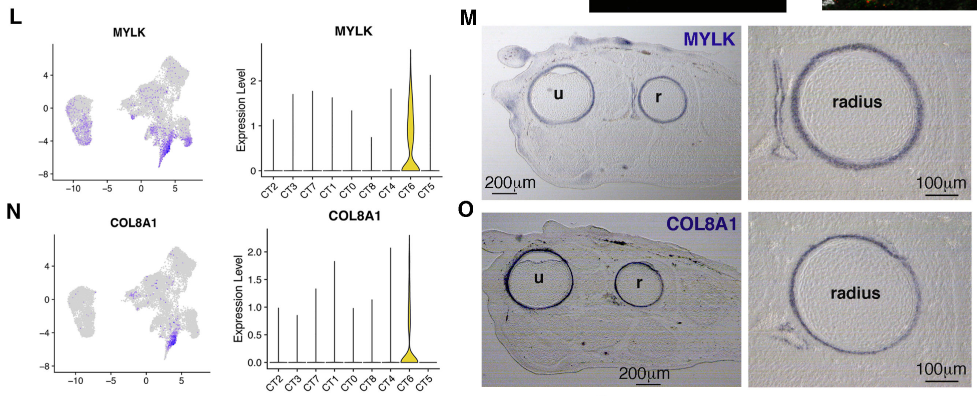


P

| | Cell number in cluster 6 | % vs cluster 6 |
|---|---|---|
| Cluster 6 | 1094 | 100% |
| MYLK+ cells | 592 | 54% |
| COL81A1+ cells | 321 | 29% |
| NET1+ cells | 488 | 45% |
| Any combinations MYLK/COL8A1/NET1 | 824 | 75% |

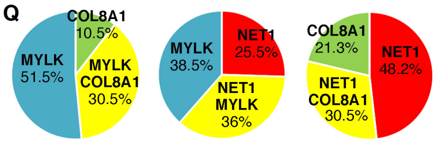


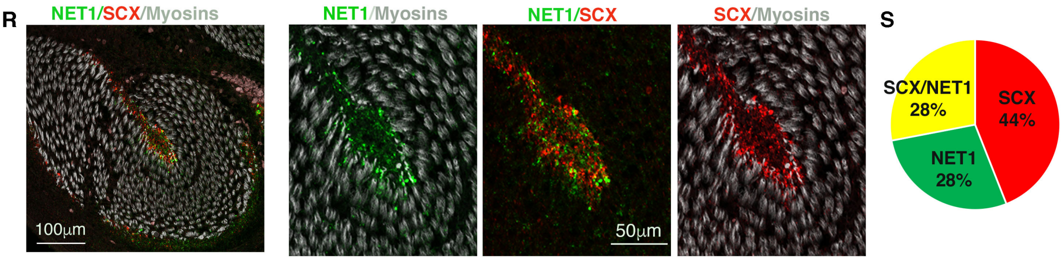


**Figure S7**

**Figure S7**

**Expression of markers for tendon fibroblast clusters**

(**A,B,C**) Expression of representative markers for tendon clusters 4, 5 and 6, *COL12A1* (A), *PTN* (B), *TGFBI* (C) with violin plots showing the log-normalized expression levels across clusters and colorimetric *in situ* hybridization to transverse limb sections of E10 chicken embryos. (**D,E,F**) High magnification of an anterior muscle of (A-C) limb sections with COL12A1 (D), PTN (E) and TGFBI (F) probes. (**G**) Adjacent section hybridized with SCX probe is shown for comparison. (**H**) Cell numbers and percentage for *COL12A1+, PTN+, TGFBI+* fibroblasts in clusters 4, 5, 6 combined.

(**I**) Cell numbers and percentage for *COL12A1+, PTN+, TGFBI+*, *ADGRG2+* fibroblasts in cluster 4. (**J,K**) Double *in situ* hybridization with ADGRG2 (cluster 4, red) and ANXA2 (cluster 1, green) probes with myosin immunostaining (grey), focused on the FCU muscle. (K) is a high magnification of part of (J). White arrows in (K) point to ADGRG2/ANXA2 expression surrounding tendon (not labeled, white stars).

(**L**) Feature plot showing the distribution of *MYLK+* cells and violin plot showing the log-normalized expression levels across clusters for the *MYLK* gene at E10. (**M**) Colorimetric *in situ* hybridization to transverse limb sections of E10 chicken embryos with MYLK probe. Right panel is a high magnification of perichondrium around the radius. (**N**) Feature plot showing the distribution of *COL8A1+* cells and violin plot showing the log-normalized expression levels across clusters for the *COL8A1* gene. (**O**) Colorimetric *in situ* hybridization to transverse limb sections of E10 chicken embryos with COL8A1 probe. Right panel is a high magnification of perichondrium around the radius. (**P**) Cell numbers and percentage of *MYKL+, COL8A1+, NET1+* cells in cluster 6. **(Q)** Percentage of *MYLK+, COL8A1+* and double *MYLK +/COL8A1+* fibroblasts among the *MYLK-COL8A1* population within cluster 6. Percentage of *MYLK+, NET1+* and double *MYLK+/NET1+* fibroblasts among the *MYLK-NET1* population within cluster 6. Percentage of *COL8A1+, NET1+* and double *COL8A1+/NET1+* fibroblasts among the *COL8A1-NET1* population within cluster 6.

(**R**) Double *in situ* hybridization with NET1 (green) and SCX (red) probes with myosins staining (grey), focused on the FCU muscle. (**S**) Percentage of *SCX+, NET1+* and double *SCX+/NET1+* fibroblasts among the *SCX-NET1* population within cluster 5. Scale bars are on *in situ* hybridization panels. u, ulna; r, radius.

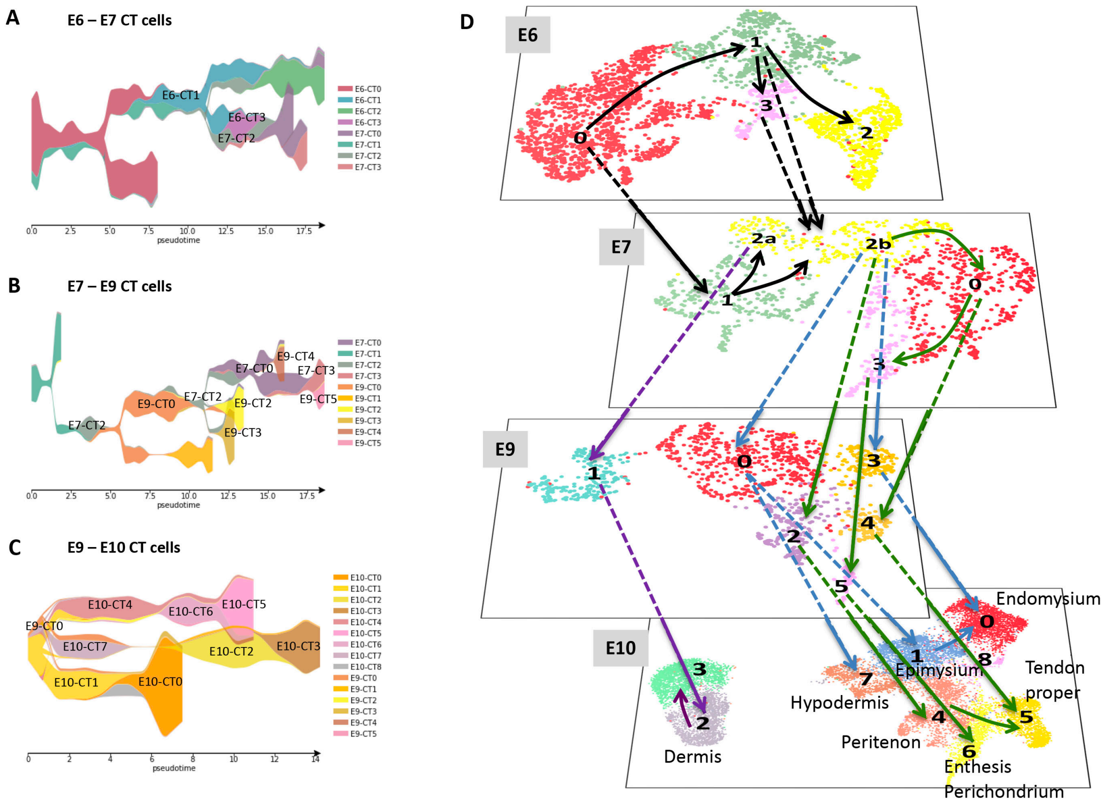


**Figure S8**

**STREAM trajectories on CT datasets of two consecutive stages combined**

**(A-C)** Dendrograms showing the cluster distribution for the E6-E7 (A), E7-E9 (B) and E9-E10 (C) CT datasets. (**D**) Representation of the CT cluster lineage tree derived from the analysis of STREAM trajectories performed on the E6-E7, E7-E9 and E9-E10 combined CT datasets. UMAP plots showing the distribution of CT clusters for the E4, E6, E7, E9 and E10 CT datasets. This schematic includes the full complement of trajectories. Black arrows refer to trajectories before lineage segregation. Blue arrows refer to MCT-related trajectories. Green arrows refer to Tendon-related trajectories. Purple arrows refer to Dermis trajectories. The cluster identity is indicated when identified.

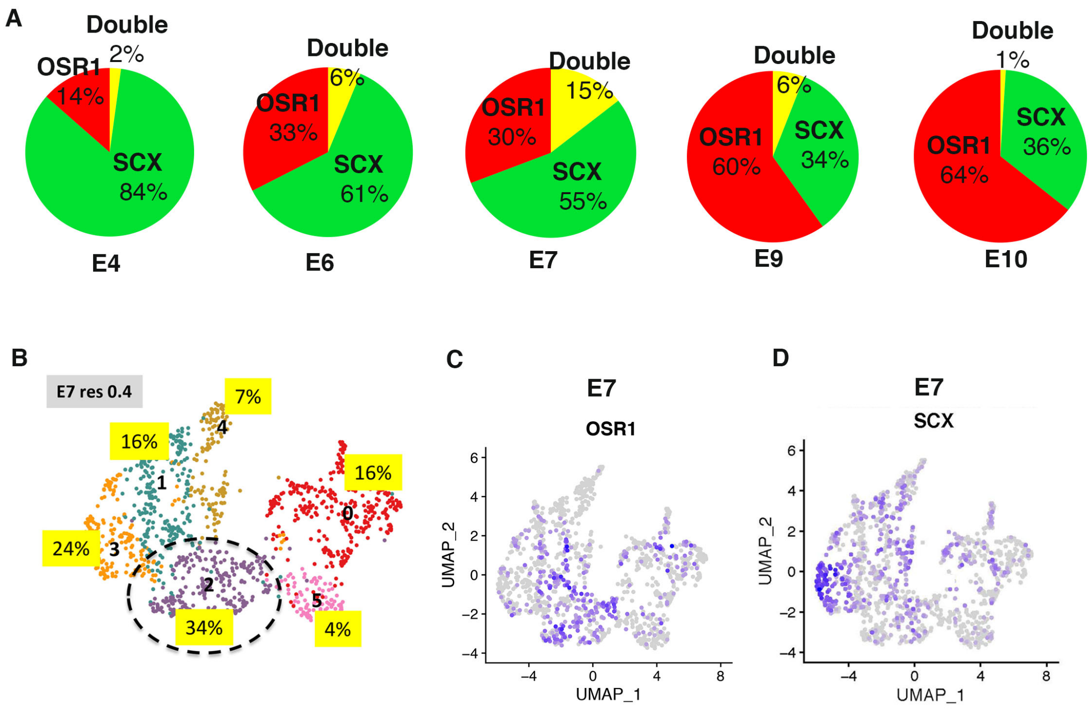


**Figure S9**

**A common population for the MCT and Tendon lineages**

(**A**) Percentage of *SCX*+ cells (green), *OSR1*+ (red) and double *SCX+/OSR1+* cells (yellow) among the SCX-OSR1 population at E4, E6, E7, E9 and E10. **(B)** UMAP plot showing the distribution of the clusters (resolution 0.4) at E7. Assigned to each cluster, the percentage of double *SCX+/OSR1+* cells of this cluster among the double SCX+/OSR1+ cells in all clusters at E7. (**C**) Feature plot showing the distribution of *OSR1*+ cells at E7. (**D)** Feature plot showing the distribution of *SCX*+ cells at E7.
