## Supplementary material for "Limb connective tissue is organized in a continuum of promiscuous fibroblast identities during development": Table S4

**Table 4**

Primer list for PCR probes

| **Gene name** | **Forward** | **Reverse** |
| --- | --- | --- |
| ADGRG2 | TGGTTTCCGTCCTCTCATTC | TGCAACAGAGCTGGTGGTAG |
| AHNAK2 | CCCAACGGGTGAGATTTCTA | ATTTGCGCCCCTTCTTCTAT |
| ALDH1A2 | TGATCCAAGAAGCAGCTGGA | GAAGAGACTGTCAGGGCCTT |
| ANXA2 | AAGGGCAAAAGGTGTGAAGA | GTCCTCTCCACCACACAGGT |
| BCL11b | GGAGGGGACAGTCAATGGAA | CATTTGTAAGGCCGCTCTCC |
| CCN3 | TGGCTGCATACAGACAGGAG | ATGGGATCTAATGGCTGGAA |
| CD34 | TCATCAGTCAGCCCCATCTC | TGAGCATCCCCTTCGCATAT |
| CHODL | CGTTCTCAGCGGTCAAAAG | GACATTGTCTGGGCCATACTT |
| COL8A1 | GTCCACTCAGGGTGGTGTCT | CTATCCCTGGGAGTCCATGA |
| COL12A1 | TCCAGCTCTTCCAAATGCTT | ACTGCTCGCATCATGTTCTG |
| EPHA3 | GTTGGAGCAGGGGAATTTGG | CTGGCTGATGTGAACTTCCG |
| HIC 1 | GGTCATCCAGGCTTGCTACT | CTCCCTCTCCAGCTCCTTCT |
| HTRA1 | AGAACAGGGTGAAGGTGGAG | GCGATCTTTCAGCTCTCCAG |
| MYLK | AACCGAGATCAGTGACAGCA | TTGGTGCTGACTTTCTTGCC |
| NET1 | TGCCCAGAAGTCATGTAGCA | TGTCCAAGAAGCTCCAGAGG |
| PDGFRA | CCTGACGCATCAGTGAAAGA | ACCATTCCACATCAGGAAGC |
| POSTN | GATCATGGGTGGAGCTGTCT | TCTGCTGGCTTGATGATTTG |
| SST | CCTGCTCTCCATCGCCTT | CCAGCTCCAGTCTCACTTCA |
| TCF7L2 | GGCACAGCTGTTTGGTCTAG | TTGTCCTCGTCCAGCTTCTT |
| TGFBI | GATCCCACGAGAGATGCTGA | TCACCCAGCGTCTGTAGTTT |
| TWIST2 | TTCTCCTGTTTCCCCTGTGG | GTCCATCTCGTCGCTCTGTA |
| ZFHX4 | CCGTGGTTTATGGAGTGCTT | AAACAGTCGCAGAGGAGGAA |
